# Design of SARS-CoV-2 RBD Immunogens to Focus Immune Responses Towards Conserved Coronavirus Epitopes

**DOI:** 10.1101/2025.01.09.632180

**Authors:** Caitlin Harris, A. Brenda Kapingidza, James E. San, Jayani Christopher, Tyler Gavitt, Brianna Rhodes, Katarzyna Janowska, Christopher O’Donnell, Jared Lindenberger, Xiao Huang, Salam Sammour, Madison Berry, Maggie Barr, Rob Parks, Amanda Newman, Mary Overton, Thomas Oguin, Priyamvada Acharya, Barton F. Haynes, Kevin O. Saunders, Kevin Wiehe, Mihai L. Azoitei

## Abstract

SARS-CoV-2 continues to evolve, with new variants emerging that evade pre-existing immunity and limit the efficacy of existing vaccines. One approach towards developing superior, variant-proof vaccines is to engineer immunogens that preferentially elicit antibodies with broad cross-reactivity against SARS-CoV-2 and its variants by targeting conserved epitopes on spike. The inner and outer faces of the Receptor Binding Domain (RBD) are two such conserved regions targeted by antibodies that recognize diverse human and animal coronaviruses. To promote the elicitation of such antibodies by vaccination, we engineered “resurfaced” RBD immunogens that contained mutations at exposed RBD residues outside the target epitopes. In the context of pre-existing immunity, these vaccine candidates aim to disfavor the elicitation of strain-specific antibodies against the immunodominant Receptor Binding Motif (RBM) while boosting the induction of inner and outer face antibodies. The engineered resurfaced RBD immunogens were stable, lacked binding to monoclonal antibodies with limited breadth, and maintained strong interactions with target broadly neutralizing antibodies. When used as vaccines, they limited humoral responses against the RBM as intended. Multimerization on nanoparticles further increased the immunogenicity of the resurfaced RBDs immunogens, thus supporting resurfacing as a promising immunogen design approach to rationally shift natural immune responses to develop more protective vaccines.

## Introduction

The rapid development of vaccines was instrumental in controlling the COVID-19 pandemic and reducing the health risk posed by SARS-CoV-2. The currently licensed SARS-CoV-2 vaccines in the United States deliver the viral spike protein formulated as mRNA or protein. First-generation vaccines contained the spike of the Wuhan-Hu-1 viral strain and offered robust protection against both infection and serious disease by eliciting high titers of neutralizing antibodies against the Receptor Binding Domain (RBD), the spike domain responsible for human ACE2 receptor attachment and viral entry^1–4^. However, the emergence of SARS-CoV-2 variants that contain a large number of mutations in the RBD caused a significant decrease in the efficacy of the ancestral vaccines^5–8^. To better protect against viruses from the omicron lineage, COVID-19 vaccines have been updated several times to include spikes from the BA.4/BA.5, XBB or JN.1 lineages. As SARS-CoV-2 continues to evolve, new variants have already emerged that escape the immune responses elicited by updated vaccines^9–11^. To address this, multiple current efforts are focused on developing vaccines that offer broad protection against existing and future coronaviruses by targeting conserved spike epitopes^12–14^. A critical factor for the success of these approaches is the ability to manipulate the natural immunogenicity of spike and focus antibody responses toward conserved, but typically subdominant, epitopes.

The RBD is the major target of humoral responses induced by SARS-CoV-2 infection or vaccination^15,16^. Multiple classes of antibodies have been isolated that target different epitopes on RBD^17–20^. Initial efforts were focused on developing antibody therapeutics that bind to the ACE2 recognition site, also called the receptor binding motif (RBM), due to their high neutralization potency^21–23^. However, the majority of such antibodies, categorized as class I and class II, lost their ability to neutralize Omicron and more recent variants due to the high number of mutations accumulated in the RBM^5,24,25^. RBD antibodies with broader cross-reactivity bind to conserved sites outside the RBM, such as the inner and outer face. Antibodies like DH1047^26^, S2X259^21^, or 10-40^27^, recognize an epitope on the RBD inner face that partially overlaps with the ACE2 binding site and block receptor binding, while those like S309^28^, or sp1-77^29^, bind to the outer face and neutralize the virus without blocking hACE2 attachment. Both inner and outer face antibodies have significantly broader recognition compared to RBM-focused monoclonal antibodies and can neutralize SARS-CoV-2 Omicron lineage viruses as well as other human or animal sarbecoviruses. While most of these antibodies have decreased affinity for SARS-CoV-2 XBB and later variants^30^, some, like S309, maintain their ability to protect against these currently circulating VOCs in animal models^31^. Therefore, due to their conservation and the breadth of the antibodies that typically target them, both the RBD inner and outer faces are promising targets for the induction of antibodies that can protect against diverse coronaviruses.

Most of the population currently has some immunity to SARS-CoV-2 acquired either through infection, vaccination, or a combination of both. This background immunity is diverse, reflecting exposure to different variants and immunizations with vaccines of different compositions and formulations. To develop next-generation coronavirus vaccines that better protect against existing and emerging viruses, novel antigens will need to focus immune responses onto conserved epitopes by overcoming the immunodominance of non-conserved spike epitopes in the context of pre-existing SARS-CoV-2 immunity. To achieve this, here we engineered and characterized “resurfaced” SARS-CoV-2 RBD-based immunogens that aim to shift immune responses away from the immunodominant RBM and onto the conserved inner and outer face regions. “Resurfaced” antigens were designed by mutating exposed SARS-CoV-2 RBD residues located outside the inner and outer face epitopes, such that the only sequence similarity between these molecules and the WT spike is in the target epitope regions. Because of this sequence overlap, resurfaced immunogens aim to preferentially stimulate antibodies with broad activity against the inner or outer face like S2X259 or S309 when used to boost immune responses elicited by the WT spike. Engineered resurfaced RBD molecules had >10% of the WT RBD residues mutated, were stable *in vitro,* and preferentially engaged target antibodies over RBM-directed ones. Immunization studies in mice revealed that they altered the balance of elicited antibodies against the RBM and the inner and outer face epitopes. Our results contribute to our understanding of SARS-CoV-2 spike immunogenicity and propose ways to rationally alter it towards the development of next-generation coronavirus vaccines with broader protection.

## Results

Antibodies against the RBD of SARS-CoV-2 can be divided into separate classes based on their epitope location and key residue contacts^20^. Class I and class II antibodies bind to the RBM (Fig. 1a), block ACE2 binding, and potently neutralize the ancestral SARS-CoV-2 virus^5,22^. These antibodies can be encoded by public clonotypes^32,33^ and they do not require extensive somatic mutations to achieve their function, which made them attractive therapeutic targets early in the pandemic. However, with the emergence of Omicron lineages containing highly mutated RBDs, many of the RBM-targeting class I and class II antibodies lost their activity^5,30,34^. Other antibody classes bind to different RBD epitopes distinct from or only partially overlap the RBM^5,35^. While their neutralization potency can be more limited compared to class I and II antibodies, their epitopes are typically more conserved resulting in broader breadth^36^. To systematically identify sub-genus level, conserved RBD regions that could be targeted by “variant-proof” vaccines, we performed sequence analysis on a representative set of RBDs selected from all known sarbecoviruses, including SARS-CoV-2 and its variants. Our results show that relative to the RBM, the RBD inner and outer face were highly conserved, with the inner face being more conserved than the outer face (Fig. 1b). This was further confirmed by comparing the median sequence conservation scores of the residues bound by ACE2 and those from the epitopes of antibodies S309 and S2X259, two canonical outer and inner face antibodies, respectively (Fig. 1b-d). The sequence conservation of the ACE2 binding footprint was significantly lower than that of S309 and S2X259. These results suggest that antibodies targeting these regions of the RBD are likely to recognize a broader set of sarbecoviruses, when compared to RBM antibodies.

**Figure 1.**
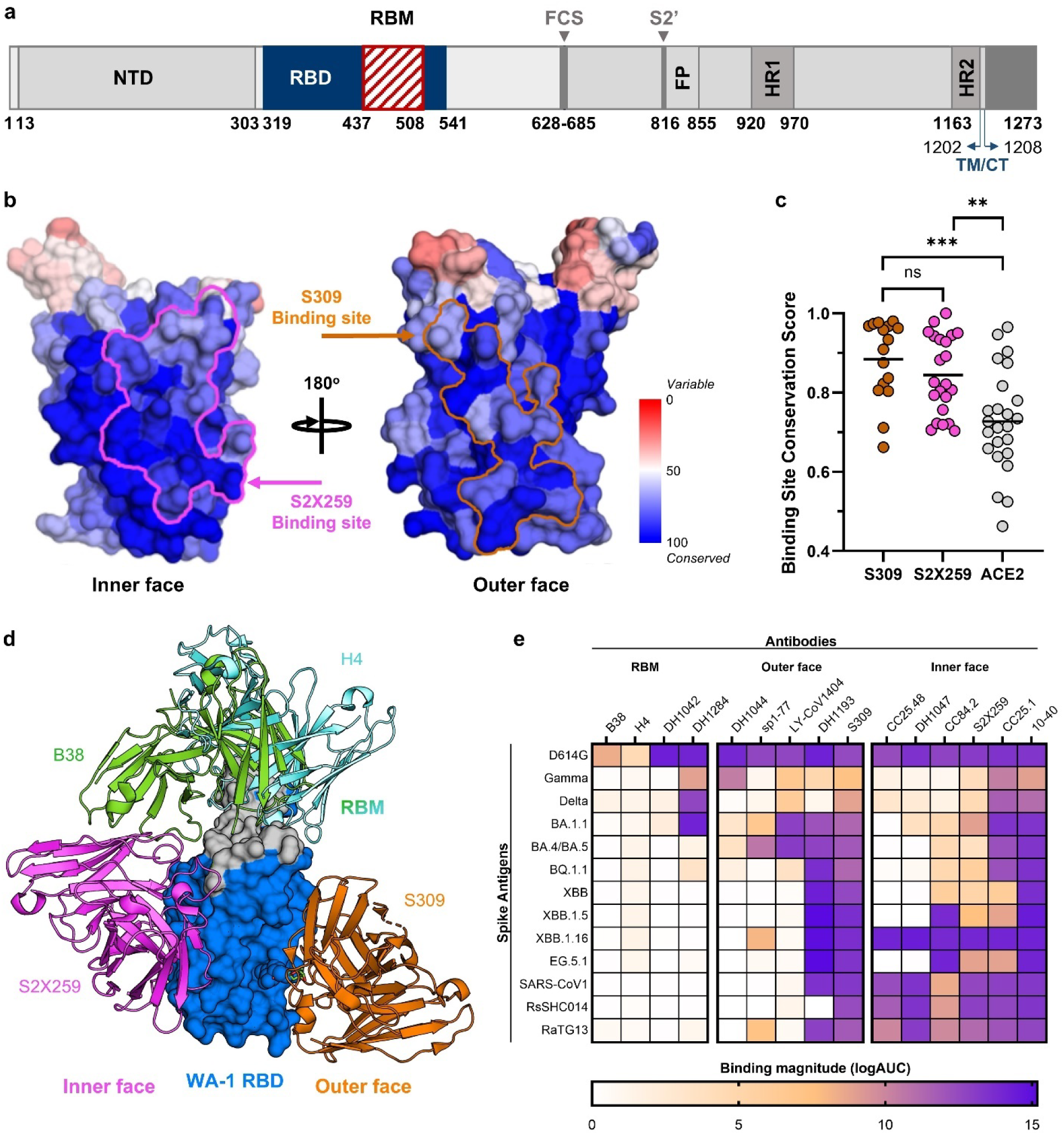
Conservation and antigenicity of the RBD across SARS-CoV-2 VoCs and other betacoronaviruses. a) SARS-CoV-2 spike protein domain organization. NTD: N-terminal domain; RBD: receptor-binding domain; RBM: receptor-binding motif; FCS: furin cleavage site; S2’: S2’ protease cleavage site; FP: fusion peptide; HR1 and HR2: heptad repeats 1 and 2; TM/CT, transmembrane/cytoplasmic tail. b) Mapping of sarbecovirus amino acid sequence conservation on the RBD structure, with the binding footprints of the inner face antibody S2X259 (*pink*) and the outer face antibody S309 (*orange*) outlined. c) Binding site conservation scores across the individual RBD amino acids involved in interactions with S309 mAb (*orange*), S2X259 mAb (*pink*), and ACE2 (*grey*), respectively. Group mean values are shown in black. *^ns^p = not significant, ***p ≤ 0.001, **p ≤ 0.01,* as calculated with Wilcoxon rank sum test. d) Structural mapping of a representative set of RBM antibodies: H4 and B38 (*cyan/green*), inner face mAb S2X259 (*pink*), and outer face mAb S309 (*orange*) onto the SARS-CoV-2 RBD. e) ELISA binding of diverse CoVs spikes to antibodies that target the RBM, outer, and inner face regions of the RBD. Values are calculated as the logarithm of the area under the ELISA binding curve.

Next, we measured, by enzyme-linked immunosorbent assay (ELISA), the binding of representative RBD antibodies targeting epitopes on the RBM, the inner or outer face, to a panel of spike proteins spanning variants from WA-1 to EG.5, and that also included proteins from SARS-CoV-2 and two pre-emergent animal viruses (Fig. 1e; Fig. S1, 2). The RBM-directed antibodies had strong binding to WA-1 but lost their reactivity to VOCs, including earlier ones like Gamma or Delta. In contrast, some inner and outer face monoclonal antibodies displayed strong binding to all the spikes tested, since their epitopes remained conserved as the RBM accumulated mutations.

We next aimed to design RBD based immunogens that preferentially elicit mAbs with broad coronavirus recognition against the conserved inner and outer face epitopes, while minimizing the elicitation of RBM-directed responses with limited breadth. To achieve this, we used a “resurfacing” strategy, where RBD surface residues outside the target epitopes were mutated. We hypothesized that by preserving the WT identity of RBD residues located in the inner and outer face epitopes, the resulting “resurfaced” immunogens will preferentially amplify immune responses against these epitopes when used to boost antibodies elicited by WT RBD exposure.

Solvent accessible sites on the RBD were identified computationally using the GETAREA web server^37–39^ by setting the side chain exposure threshold above 40%. Next, surface residues were mutated, except for prolines or hydrophobic residues that were judged to pack against other residues upon structural analysis. Initially, mutations were informed by computational modeling with Rosetta^40,41^. Although selected mutations were predicted to be favorable, the majority of RBD variants engineered with this approach tested experimentally were expressed poorly or unstable. The choice of mutations at “resurfaced” sites was greatly facilitated by the deep scanning mutagenesis analysis of the Wuhan RBD that revealed the effect of every amino acid substitution on expression level and receptor binding^42^. Hence, mutations at a given position were selected from characterized substitutions that change the chemical character of the original amino acid but were shown to maintain RBD expression. However, even with this approach, we found that combinations of mutations predicted to be favorable for expression often resulted in constructs that had low yields and stability, likely due to unknown interference effects.

The first successful design we characterized, named Resurf6, contained 22 mutations relative to the WT RBD, about ∼10% of the total residues, with four changes located in the RBM and 18 positioned outside (Fig. 2a, b). The RBM was mutated at “hotspot” residues for ACE2 binding that were also engaged by RBM antibodies (Y449, L455, Q493, and Y505). The inner and outer face epitopes of S2X259 and S309 were kept unchanged, except for one mutation located at the periphery of the S309 epitope (L441Q) that was expected to improve expression without affecting antibody binding (Fig. 2c). Binding analysis by ELISA and SPR revealed that Resurf6 had strong binding to all the inner face antibodies tested, on par with WT RBD (Fig. 2d, e; Fig. S3, 4). Resurf6 maintained high-affinity binding of outer face antibodies S309^28,43^, LY-CoV1404 and sp1-77^29^, but lost binding to antibodies with footprints that contact both the outer domain and the periphery of the RBM like DH1193 and DH1044^44^ as expected. The binding of RBM antibodies H4 and B38^22^ was eliminated, but Resurf6 still bound the class II monoclonal DH1042, suggesting that additional modifications are necessary to limit interactions of RBM antibodies further. To achieve this, we developed a second-generation resurfaced RBD molecule, called Resurf61, which contained four additional mutations in the 481-491 loop (F486T, N487E, Y489V and F490A) (Fig. 2a-c). This loop harbors hotspot residues crucial for binding of class II antibodies like DH1042^45^. Resurf61, which contained 26 mutations relative to WT RBD, had a binding profile like Resurf6 to inner and outer face antibodies, but no longer interacted with any of the class I and II mAbs tested (Fig. 2d, e). In an alternative approach, a different set of mutations was introduced into the RBM of Resurf6 to develop another resurfaced design named Resurf64. In this construct, four new RBM changes were added to Resurf6 (K417N, K444N, G446V, E484K) while two existing Resurf6 mutations were reverted to WT to improve expression and stability (Fig. 2a-c). Resurf64 showed no binding to any of the class I and II antibodies tested by 1:1 SPR measurements, although some low-level binding was detected by ELISA, in an assay where avidity was present (Fig. 2d, e). Therefore, in addition to using a different mechanism for blocking RBM antibodies, Resurf64 only bound with high affinity to the broadest outer face antibodies like S309 to improve immunofocusing further. The resurfaced RBDs were thermodynamically stable despite having >10% of the WT RBD residues mutated (Fig. 2f). Melting temperatures were >44°C, but 5-9 degrees lower than for WT RBD (Tm = 53.7°C).

**Figure 2.**
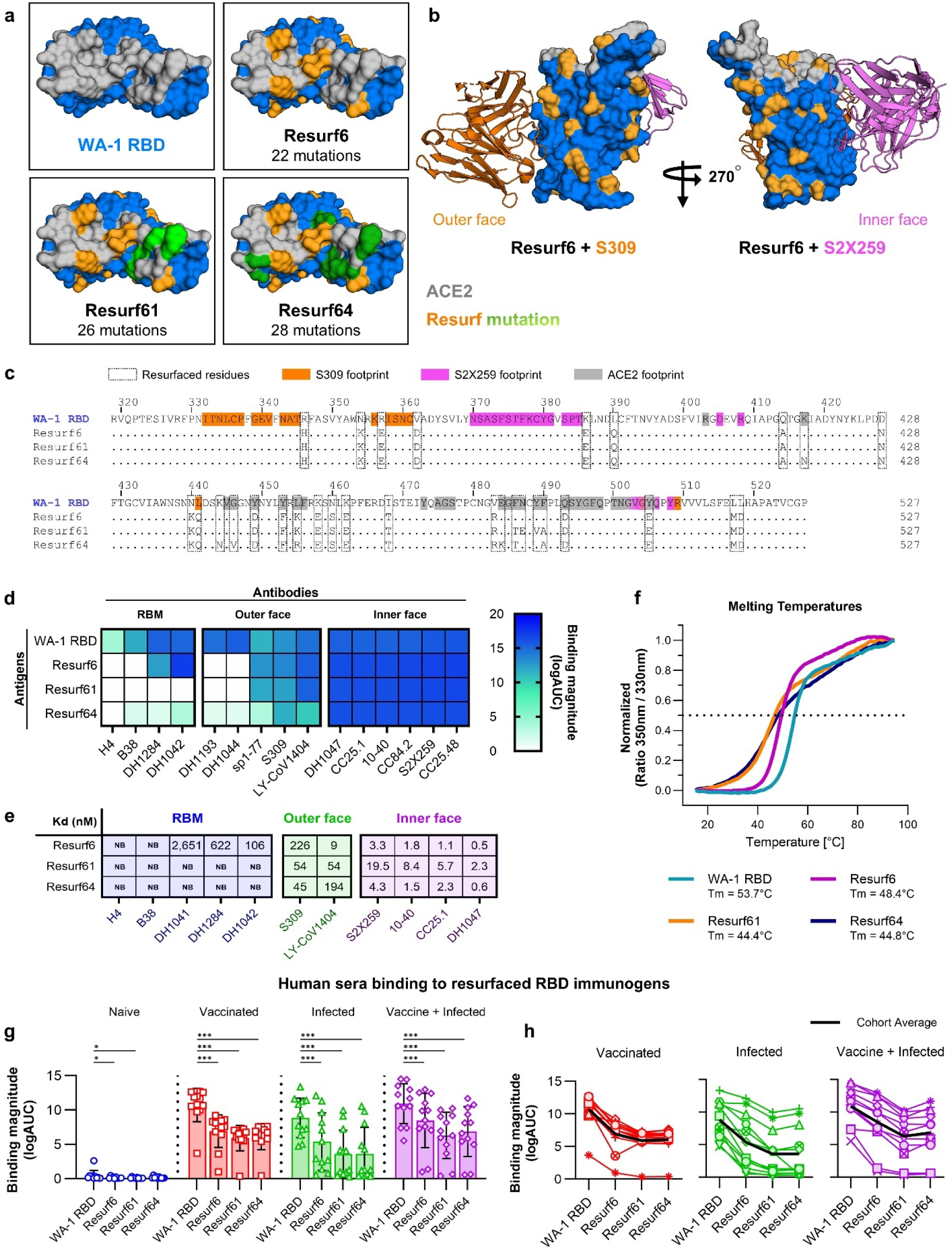
Design and *in vitro* characterization of resurfaced RBD immunogens. a) Top view of wild-type RBD and the three engineered resurfaced RBD immunogens, with mutated sites in *orange* and *green*. ACE2 binding residues are shown in *grey*. b) Side view of RBD in complex with S309 (*left*) and S2X259 (*right*) antibodies. Residues modified in Resurf6 are shown *orange*. ACE2 binding residues are shown in *grey*. c) Sequence alignment between WA-1 RBD and the three resurfaced RBD immunogens, with modified residues delineated in boxes. Residues binding to S309 mAb, S2X259, and ACE2 are colored *orange*, *pink*, and *grey*, respectively. d) Binding of WT and resurfaced RBDs to mAbs targeting diverse epitopes on RBD. Values represent the logarithm of the area under the curve. e) Binding affinities of resurfaced immunogens to RBM, outer and inner face mAbs. f) Melting temperature curves of WA-1 RBD (*teal*) and the resurfaced immunogens Resurf6 (*pink*), Resurf61 (*orange*), and Resurf64 (*blue*). g) Binding of human sera from subjects naïve (*blue*) or with existing SARS-CoV-2 immunity acquired by vaccination (*red*), infection (*green*), or vaccination followed by infection (*purple*) to WT and engineered RBDs. Data are calculated as the logarithm of the area under the ELISA binding curve and are presented as the mean +/- standard deviation. h) The same data as in (g) plotted for each subject. The cohort average is shown in *black*. For graphs (g) and (h), *n = 12* independent subjects. Significance tested using the Wilcoxon signed-rank test. *\*\*\*p ≤ 0.001 and *p ≤ 0.05*.

Next, we analyzed the binding of the resurfaced RBD immunogens to polyclonal sera obtained from subjects with pre-existing immunity to SARS-CoV-2 acquired by either infection, vaccination, or infection followed by vaccination, from a cohort we previously described^12^. Each of these three groups contained samples from 12 individuals, with the majority vaccinated with first-generation mRNA or adenovirus vaccines and infected with Delta or Omicron variants of SARS-CoV-2. As expected, sera from all three groups had strong binding to the WT WA-1 RBD (Fig. 2g, h; Fig. S5). Importantly, sera binding to the resurfaced RBDs was reduced at least two-fold, with the highest decrease observed for Resurf61 and Resurf64, the two constructs that had no binding to RBM mAbs (Fig. 2g, h). This drop was observed in all the individual samples. Therefore, the engineered resurfaced RBDs have reduced recognition of not just RBM mAbs, but also polyclonal sera, consistent with the design goal of mutating away immunodominant epitopes that are the major target of antibodies elicited by spike. Because of their antigenic characteristics, the resurfaced RBD can be used as probes to determine the ratio of sera responses binding to the immunodominant RBM relative to other more conserved epitopes. Of note, half the samples in the infection only group showed no binding to Resurf61 and Resurf64, suggesting that elicited humoral responses almost exclusively target the RBM in these individuals (Fig. 2g, h). This contrasts with the vaccination only and infection followed by vaccination groups where 11/12 and 10/12 samples respectively maintained robust, albeit two-fold reduced, binding to Resurf61 and Resurf64. This supports the observation that vaccination or hybrid immunity typically elicit broader responses relative to natural infection alone^46,47^.

Resurfaced RBD immunogens were next used to vaccinate BALB/c mice both on their own and as boosts to mRNA encoding WA-1 spike (Fig. 3a). Animals received three shots, 4 weeks apart, and sera were collected and analyzed pre-vaccination and one week after each immunization. Sera from animals immunized with Resurf6, Resurf61, and Resurf64 showed significantly reduced binding to WA-1 Spike and RBD compared to animals immunized with WT WA-1 RBD, suggesting a difference in immune responses consistent with a reduction in the RBM antibodies that typically dominate responses induced by native SARS-CoV-2 proteins (Fig. 3b; Fig. S6). Conversely, WA-1 RBD elicited sera had reduced binding to the resurfaced RBDs, with decreased binding to the antigens according to their ability to recognize RBM antibodies, highest binding to Resurf6 and lowest to Resurf64 (Fig. 3b). Interestingly, WA-1 RBD elicited sera showed robust binding to the resurfaced immunogens, suggesting the induction of antibodies against the inner and outer face epitopes upon repeat immunizations with native SARS-CoV-2 proteins (Fig. 3b). The resurfaced RBD molecules were next used as boosts in animals previously immunized with WA-1 spike as mRNA and the elicited sera was compared with that from animals that were boosted with the WA-1 RBD. Sera binding was tested on a panel of recombinant spike proteins covering representative SARS-CoV-2 variants of concern (VOCs) from WA-1 to EG5 (Fig. 3c; Fig. S7). Sera binding titers were highest to WA-1/D614G and progressively decreased to spikes from VOCs, with the lowest binding detected against XBB1.5 and EG5.1 spikes. Sera was also tested for live virus neutralization against WA-1, Delta, and XBB1.5 viruses (Fig. 3d). No neutralization activity was detected against XBB1.5, despite considerable binding titers detected by ELISA, but consistent with previous observations that the ancestral spike does not elicit cross-reactive antibodies against this strain^48–50^. Boosting with WA-1 RBD induced the highest neutralization titers against WA-1 or Delta viruses. Resurf6 either on its own or followed by Resurf64, induced the highest neutralization titers from all the resurfaced constructs. While average titers were lower than for WT RBD, the difference was not statistically significant (Fig. 3d). Taken together, the binding and neutralization data revealed no significant expansion in sera breadth in animals boosted with the resurfaced RBDs compared to the WT WA-1 RBD.

**Figure 3.**
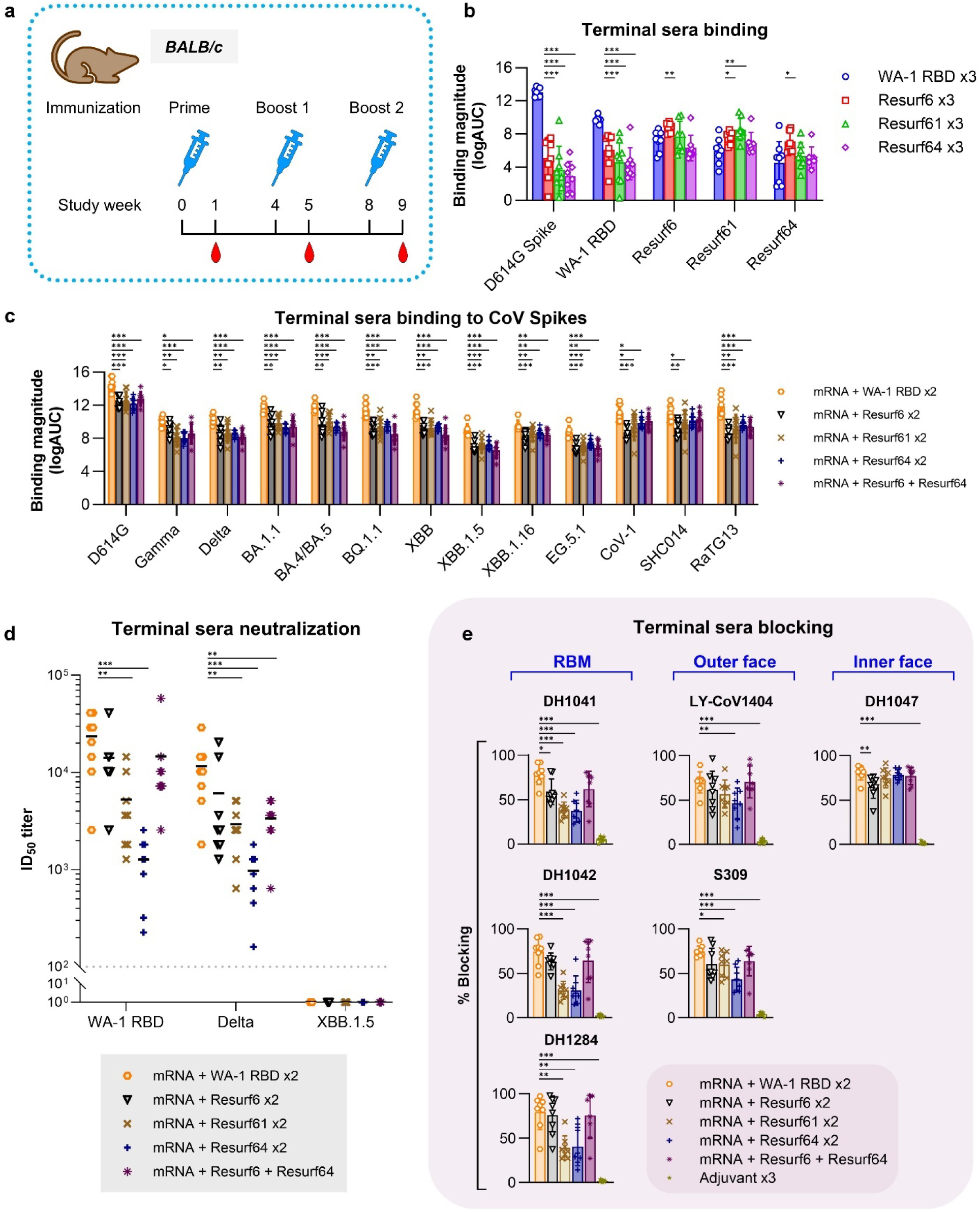
*In vivo* characterization of resurfaced RBD immunogens. a) Study design to determine the immunogenicity of monomeric resurfaced immunogens in BALB/c mice. b) Binding of terminal sera from animals immunized with different RBD immunogen against SARS-CoV-2 WA-2 spike, WA-1 RBD, and the resurfaced immunogens. Data are calculated as the logarithm of the area under the ELISA binding curve and are presented as the mean +/- standard deviation. c) Binding of terminal sera from animals primed with mRNA and boosted with different RBD immunogens against diverse CoV spikes. D614G, Gamma, Delta, BA.1.1, BA.4/BA.5, BQ.1.1, XBB, XBB.1.5, XBB.1.16 and EG.5.1 refer to the respective SARS-CoV-2 variants. CoV-1 = SARS-CoV-1; SHC014 = BatCoV RsSHC014; RaTG13 = BatCoV RaTG13. Data are calculated as the logarithm of the area under the ELISA binding curve and are presented as the mean +/- standard deviation. d) Live virus neutralization of the sera from (c). e) Blocking of terminal sera from animal groups in (c) to RBM (DH1041, DH1042 and DH1284), outer (LY-CoV1404 and S309), and inner (DH1047) face targeting mAbs. Data are presented as the mean +/- standard deviation. For graphs (b - e), all groups *n = 8* represent independent mice. Significance tested using the Wilcoxon signed-rank test. *\*\*\*p ≤ 0.001, **p ≤ 0.01, *p ≤ 0.05*.

To better characterize the specificity of the elicited sera, we performed blocking experiments, where the binding of an antibody with a well-characterized epitope on spike was measured in the presence or absence of the induced sera. Blocking of six representative antibodies binding to WA-1 spike that targeted the RBM (DH1041, DH1042, DH1284), inner face (DH1047), or outer face (S309, LY-CoV-1404) epitopes were analyzed in this fashion (Fig. 3e). For all the three RBM antibodies tested, sera samples from animals boosted with Resurf61 and Resurf64 had significantly less blocking than those from animals boosted with the WT WA-1 RBD. This indicates limited boosting of RBM-directed responses by the engineered immunogens, in line with our design goals and consistent with the *in vitro* binding of monoclonal antibodies and polyclonal humoral responses. Based on this analysis, we can infer those sera from animals boosted with Resurf61 or Resurf64 contained 2-3-fold less RBM antibodies than that elicited by boosting with WT RBD. Resurf6 boosted RBM-directed response better than the other resurfaced constructs but worse than WT RBD, consistent with its antigenic profile. Average sera blocking for outer face antibodies LY-CoV1404 and S309 was uniformly reduced in the resurfaced RBD immunized animals compared to the WT RBD group, although not all decreases were deemed statistically significant (Fig. 3e). Blocking of inner face antibody DH1047 was similar for WT RBD, Resurf61, and Resurf64, but reduced for Resurf6. Taken together, these results show that while Resurf61 and Resurf64 successfully limited the elicitation of RBM antibodies, they did not preferentially boost inner or outer face antibodies with broad cross-reactivity.

Previous work demonstrated that RBD immunogens multimerized on nanoparticles can increase the overall titers and the breadth of the elicited humoral responses^14,51^. Therefore, we developed nanoparticles containing 60 copies of the resurfaced RBDs conjugated to the mi03 nanoparticle using the SpyCatcher-SpyTag system^52^ (Fig. 4a). Particles displaying Resurf64 were unstable, thus this immunogen was dropped from subsequent analysis. Mi-03 nanoparticles displaying Resurf6 or Resurf61 were well formed by Negative Stain Electron Microscopy (NSEM) and maintained the antigenic profile of the monomeric proteins (Fig. S8). These nanoparticles were compared for their ability to boost broad responses with a previously described ferritin nanoparticle that displays 24 copies of the Wuhan/WA-1 RBD^14^. Groups of eight mice were immunized four weeks apart, first with WA-1 spike mRNA and then boosted twice with nanoparticles displaying the WA-1 RBD, Resurf6, Resurf61, or with Resurf6 followed by Resurf61 (Fig. 4b). Sera were collected one week after each immunization and analyzed for binding, neutralization, and blocking as above. The sera from animals boosted with either WT RBD or Resurf6 nanoparticles had similar breadth and binding levels against a panel of diverse CoV spikes (Fig. 4c; Fig. S9). This contrasted with the results for boosting with the respective monomeric proteins, where WT WA-1 RBD elicited significantly higher titers than Resurf6 (Fig. 3c), suggesting that nanoparticle presentation alters the immune responses. Resurf61 boosting resulted in uniformly lower titers than WT RBD of Resurf6, while the binding titers induced by the mixed boosting regimen of Resurf6 followed by Resurf61 were similar to Resurf6 alone. Sera elicited by WA-1 RBD, and Resurf6, either by itself or followed by Resurf61, were also tested for neutralization against live viruses spanning multiple SARS-Cov-2 VOCs and SARS-Cov-1 (Fig. 4d). Average titers elicited by boosting with Resurf6 followed by Resurf61 were generally higher than those induced by WT RBD, but the increase did not reach statistical significance, except against Gamma. A similar pattern was observed for boosting twice with Resurf6. Intriguingly, half the animals boosted twice with Resurf6 elicited detectable neutralization titers against XBB1.5 (Fig. 4d). Sera was also tested in a pseudovirus neutralization assay that included the JN.1 variant and bat coronavirus SHC014 (Fig. S10). No significant activity was observed against JN.1, and the overall results were in agreement between the two neutralization assays. Taken together, these results suggest that boosting with the resurfaced immunogens multimerized on nanoparticles leads to the elicitation of antibodies with increased breadth compared to boosting with the same proteins as monomers. To map the major epitopes targeted by the elicited sera, we performed blocking experiments against well characterized antibodies as above (Fig. 4e). Interestingly, sera from animals boosted with Resurf6 blocked RBM antibodies DH1041, DH1042 and DH1284 at least as well as that from animals boosted with WT RBD. This contrasts with the results obtained with the respective monomeric proteins, where RBM mAb blocking was lower for sera boosted by the resurfaced designs (Fig. 3e), suggesting that nanoparticle presentation amplifies responses against the RBM. Similarly, sera blocking against RBM class I antibody B38 was significantly higher in animals boosted with Resurf6 followed by Resurf61 than in those boosted with WT RBD (Fig. 4e). Regarding blocking of outer and inner face antibodies, elicited sera from animals boosted with Resurf6 were similar to that from the animals that received WT RBD, as also observed for the monomeric proteins; the group averages trended higher, but they were not statistically significant.

**Figure 4.**
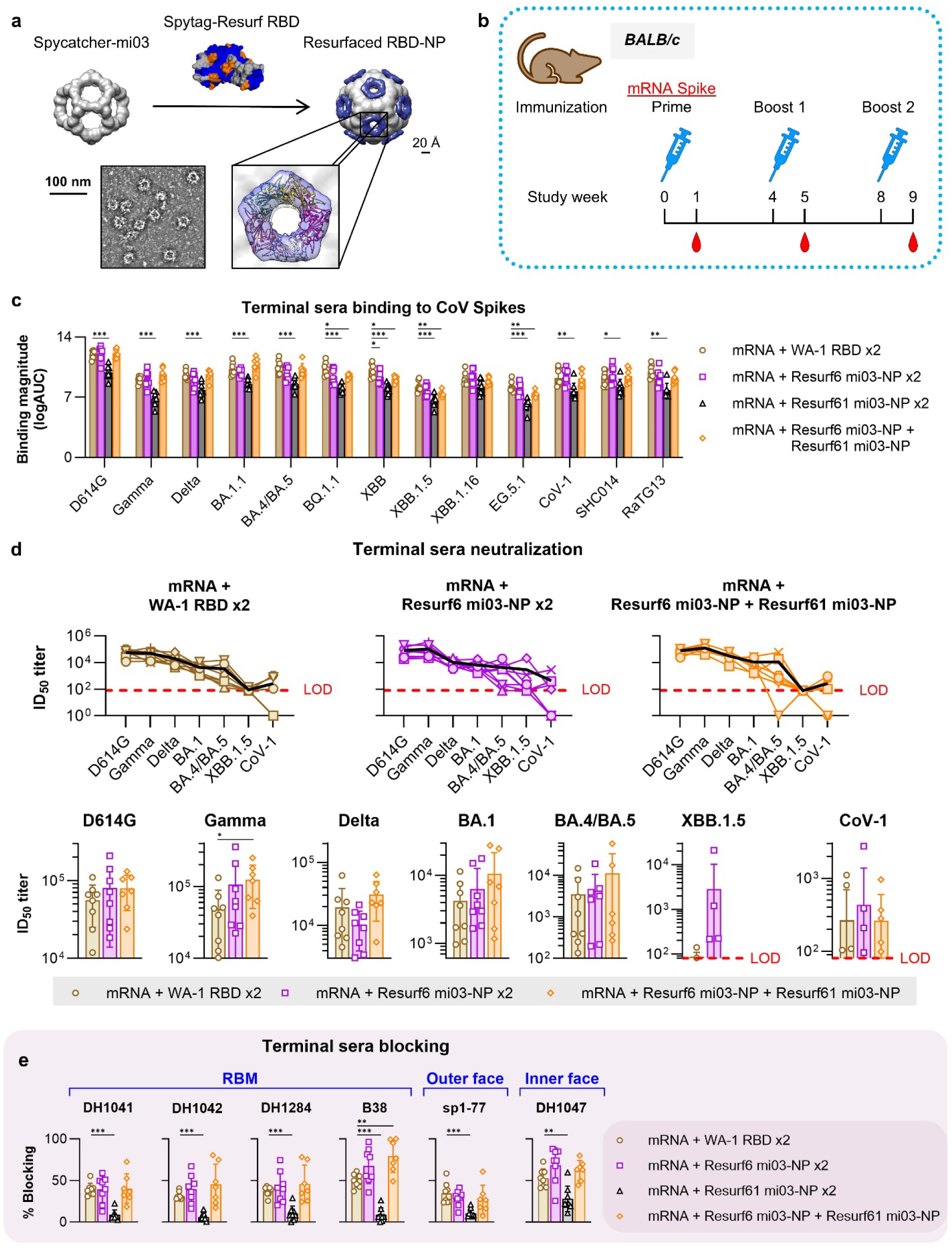
Immunogenicity of resurfaced RBD immunogens multimerized on nanoparticles. a) Development of mi03 nanoparticles (NP) displaying multimerized resurfaced RBD immunogens. b) Study design to determine the immunogenicity of multimerized NPs displaying resurfaced immunogens in BALB/c mice. c) Binding of terminal sera from animals primed with D614G mRNA spike and boosted with the RBD immunogens to diverse CoV spikes. D614G, Gamma, Delta, BA.1.1, BA.4/BA.5, BQ.1.1, XBB, XBB.1.5, XBB.1.16 and EG.5.1 refer to the SARS-CoV-2 variants. CoV-1 = SARS-CoV-1; SHC014 = BatCoV RsSHC014; RaTG13 = BatCoV RaTG13. Data are calculated as the logarithm of the area under the ELISA binding curve and are presented as the mean +/- standard deviation. d) Neutralization of terminal sera from groups in (c) to different CoV live viruses. *Top:* data plotted for individual animals in each group against the live virus panel. Group averages in *black* lines. Level of detection (LOD) in *red*. *Bottom:* average group neutralization titers against each live virus. Error bars show +/- standard deviation. e) Blocking of the terminal sera from immunizations groups in (c) to RBM (DH1041, DH1042, DH1284 and B38), outer (sp1-77) and inner (DH1047) face mAbs. Data are presented as the mean +/- standard deviation. For graphs (c - e), all groups *n = 8* represent independent mice, except for mRNA + Resurf6 mi03-NP + Resurf61 mi03-NP *(n = 7)* due to a mouse death prior to terminal bleeds. Significance tested using the Wilcoxon signed-rank test. *\*\*\*p ≤ 0.001, **p ≤ 0.01, *p ≤ 0.05*.

Taken together, these results show that, compared to monomeric protein, resurfaced RBD immunogens displayed on nanoparticles increase the titers and breadth of the elicited sera, likely due to increased antibody titers against the RBM.

## Discussion

As SARS-CoV-2 continues to circulate widely, new immune-evasive variants will continue to emerge, highlighting the need for novel vaccines. Currently, SARS-CoV-2 vaccines are updated periodically to best match the circulating viral strains, as is the case with the seasonal influenza vaccine. An alternative approach is to develop vaccines that offer broad protection against both current and future SARS-CoV-2 variants by preferentially eliciting antibodies against conserved sites of the spike protein. The RBD is the primary target of neutralizing antibodies elicited by current vaccines or natural infection. However, the RBD is highly variable, especially in the RBM^36^, allowing it to readily accumulate mutations that escape existing immunity^53^.

Our analysis and previous reports showed that the inner and outer faces of the RBD are conserved across betacoronaviruses, suggesting that antibodies targeting these conserved sites could offer protection against a broad spectrum of sarbecoviruses, including animal ones with spillover potential. Individuals exposed to or vaccinated against SARS-CoV-2 generated antibodies against the inner and outer faces of RBD, but they typically make up only a small fraction of the total humoral responses^54^. Inner and outer face antibodies show broad neutralization, effectively targeting multiple SARS-CoV-2 VoCs and other coronaviruses. For example, the monoclonal antibody S309 neutralized SARS-CoV-1, SARS-CoV-2 and SARS-CoV-2 VOCs, including Omicron sublineages BA.1, BA.1.1 and BA.2 and Beta^28,55^. However, these antibodies can be susceptible to escape mutations. For example, the S309 antibody failed to bind a SARS-CoV-2 strain containing the P337L mutation^56^. Nevertheless, the conservation of the inner and outer faces of the RBD within sarbecoviruses, along with the accessibility of the outer face in both open and closed RBD states, suggests that these epitopes are promising targets for a broadly protective pan-sarbecovirus vaccines. It is likely that additional, yet undiscovered, antibodies with broad cross-reactivity may exist against the inner and outer face epitopes and that a polyclonal response containing such antibodies that interact with the target epitopes in different ways may minimize the risk of immune escape and thus provide improved breadth beyond the existing mAbs.

The goal of the resurfaced RBD immunogens was to preferentially activate and expand existing B cells, primed by exposure to the WT spike protein, which can secrete antibodies against the inner and outer face regions. To achieve this, the immunodominant RBM and other exposed amino acids outside the target epitope were mutated based on structural, computational, and deep sequence mutagenesis analysis. This approach resulted in stable immunogens that contained >10% of their amino acids mutated compared to WT RBD. Extensive *in vitro* antigenic characterization showed that the engineered RBD molecules preferentially bound to antibodies that target the inner and outer face but not the RBM, highlighting the success of our rational resurfacing approach. This approach can be generalized to the design of other immunogens and combined with complementary approaches, such as glycosylation^57^ or nanoparticle presentation^14,58^, to minimize immune responses against undesired epitopes. The design of the resurfaced RBD was greatly aided by the existence of deep scanning mutagenesis data that measured the effect of each RBD mutation on the stability of the protein^42^. However, this data set lacks information on the interference effects of multiple mutations, which affected the design of the resurfaced RBDs; combinations of single mutations that did not affect stability by deep scanning mutagenesis often led to constructs with decreased expression and poor stability. Recently, machine learning protein engineering approaches like ProteinMPNN^59^ proved adept at generating sequences that fold onto a target backbone structure. Indeed, ProteinMPNN was previously applied successfully to resurface vaccine candidates for increased solubility^12^. The design of the resurfaced RBDs also illustrates the remarkable plasticity of the SARS-CoV-2 RBD. Our results demonstrate that the structural fold of the SARS-CoV-2 RBD can accommodate a large number of mutations, suggesting that the virus could continue to evolve with a limited loss of fitness.

When used *in vivo* in animals with pre-existing spike immunity, the resurfaced RBD immunogens greatly reduced the RBM titers, per the immunogen design goals. However, no significant titer increase was observed for inner and outer face RBD antibodies. This outcome could be due to multiple factors that will be explored in future studies. It has been recently shown that the reentry of B cells generated by priming immunogens into second germinal centers is limited in mice, which could minimize the boosting effect of our resurfaced RBD immunogens^60^. However, other studies found higher levels of memory B cell reactivation after a boost, especially when the repeat immunizations are performed ipsilaterally^61^, i.e., in the same leg, as was done here. Some of the RBD inner and outer face antibodies have long CDRH3 loops of over 20 amino acids, which are rare in the mouse immune repertoire^62^, meaning that other *in vivo* models may be more appropriate to test our immunogens. It is also possible that the resurfacing mutations introduced new immunogenic sites that competed with the inner and outer epitopes, even as the RBM immunodominance was reduced. To better understand the boosting capacity of the resurfaced RBDs, it will thus be necessary to perform detailed B cell receptor repertoire analyses in the germinal centers, memory, and plasma compartments. A previous study showed that non-human primates have the capacity to elicit antibodies against the inner face of RBD^14,63^, so this animal model may be more appropriate for testing resurfaced RBD immunogens.

Interestingly, presenting the resurfaced RBDs as multimers on nanoparticles altered the boosted immune response compared to the monomers by increasing the titers against the RBM. This was accompanied by higher sera binding and neutralization breadth against diverse VOCs, suggesting that the RBDs displayed on nanoparticles adopt an orientation that favors broader responses against the RBM. The RBM antibodies elicited by the nanoparticles are likely different than the class I and II ones, which were disfavored by the designed mutations. While this hypothesis cannot be confirmed without detailed immune repertoire analysis and antibody isolation, some isolated antibodies against the RBM, like S2K416^64^, are reported to be remarkably broad and bind epitopes present on Resurf6. Taken together, the resurfaced immunogens represent a new, rational approach to disrupt the natural immune responses against SARS-CoV-2 that typically target the RBM and result in strain-specific antibodies that can be readily escaped by viral evolution. Resurfaced RBD immunogens minimize RBM directed responses under conditions of pre-existing immunity currently found in the population, and the breadth of the antibodies they elicit may be improved by nanoparticle presentation. The resurfaced RBDs favor interactions with antibodies against the RBD inner and outer faces and could contribute to vaccination regimens that favor the elicitations of antibodies against these conserved epitopes.

## Methods

### Protein Expression and purification

#### Resurfaced RBD immunogens

Resurfaced RBDs were expressed using the optimized SARS-CoV-2 RBDs expression by Stadlbauer, et al^65^. Briefly, 100 mL cultures of Expi293F cells at a density of 3.0×10^6^ cells/mL were transiently transfected with resurfaced RBDs encoding plasmids and Expifectamine (Invitrogen). 72-hours after transfection, cells were separated from culture media by centrifugation, and the supernatant was filtered with 0.2-micron filters. The supernatant was incubated with Ni-NTA beads (Qiagen) equilibrated with native binding buffer (50 mM NaH2PO4, 500 M NaCl, 10 mM imidazole, pH 7.5) for an hour at 4°C. The beads were settled by centrifugation and the supernatant was removed by pipetting. The beads were washed with wash buffer (50 mM NaH2PO4, 500 M NaCl, 30 mM imidazole, pH 7.5) after which the protein was eluted with elution buffer (50 mM NaH2PO4, 500 M NaCl, 250 mM imidazole, pH 7.5). Protein expression and purity was confirmed by SDS-PAGE analysis and quantified spectro-photochemically at 280 nm using Nanodrop 2000 (Thermofisher).

#### WT spikes and RBD immunogens

SARS-CoV-2 spike proteins were prepared using methods described previously^66^. Plasmids encoding the SARS-CoV-2 spike ectodomains were transfected into Gibco FreeStyle 293F cells following the manufacturer’s recommendations. Six days post-transfection the supernatant was harvested and filtered through a 0.22 µm filter. The spike proteins were purified via their C-terminal TwinStrep tag using StrepTactin resin (IBA LifeSciences), followed by size exclusion chromatography (SEC) on Superose 6 10/300 GL increase column (Cytiva, MA) in 2 mM Tris, pH 8.0, 200 mM NaCl, 0.02% NaN3. The quality of the protein preparations was checked by SDS-PAGE. The final products were flash-frozen in liquid nitrogen and stored at −80°C in single-use aliquots for future use. Aliquots were thawed at 37°C for 20 minutes before use.

SARS-CoV-2 RBD were purified as described previously^44^. Plasmids encoding the RBD were transfected in 293F cells and harvested from supernatant on the 6th day post transfection. All purification steps were performed at room temperature. The RBDs contained a C-terminal 6x-Histidine tag and were purified via nickel affinity chromatography using a HisTrap excel column (Cytiva, MA). The supernatant was loaded onto the column and the column washed with buffer A (1x PBS pH 8.0) until baseline. A gradient over 20 CV from 100% buffer A to 100% buffer B (1x PBS pH 8.0, 1 M Imidazole) was applied to elute the proteins from the column. Fractions containing RBD were pooled, concentrated, and further purified by size exclusion chromatography (SEC) using a Superdex 200 Increase 10/300 GL column (Cytiva, MA) equilibrated with 1x PBS, pH 8.0. Protein quality was assessed by SDS-PAGE using NuPage 4-12% (Invitrogen, CA). The purified proteins were flash frozen and stored at −80°C in single-use aliquots. Each aliquot was thawed at 4°C before use.

#### Antibodies

Antibodies were produced as we described previously^12^. Briefly, Expi293F cells at a density of 2.5×10^6^ cells/mL were transiently transfected with an equimolar plasmid mixture of heavy and light chain using Expifectamine (Invitrogen). Five days post transfection, cells were separated from culture media by centrifugation and the supernatant was filtered with 0.8-micron filter. Supernatant was incubated with equilibrated Protein A beads (ThermoFisher) for an hour at 4°C. Beads were washed with 20 mM Tris, 350 mM NaCl at pH 7 and antibodies were eluted with a 2.5% Glacial Acetic Acid elution buffer and buffer exchanged into 25 mM Citric Acid, 125 mM NaCl buffer at pH 6. Antibody expression and purity was confirmed by SDS-PAGE analysis and quantified by measuring absorbance at 280 nm (Nanodrop 2000).

### Thermal Stability

Nano Differential Scanning Fluorometry (nanoDSF) assays were performed to determine melting temperatures of the proteins using Prometheus Panta (NanoTemper Technologies). All proteins were formulated in Phosphate Buffer (20 mM sodium phosphate monobasic, 150 mM NaCl, pH 7.4). Intrinsic fluorescence was measured at 330 nm and 350 nm while the sample was heated from 15 to 95°C at a rate of 1°C /minute. The ratio of fluorescence (350/330 nm) and inflection temperatures (T) were calculated by the PR. Panta Analysis v1.6.3 software. The data analysis and the thermal denaturation curves were plotted in GraphPad Prism 9.5.1.

### Antigenicity

#### Surface Plasmon Resonance

The dissociation constants (*K*_D_s) between the resurfaced RBDs and target antibodies were determined by Surface Plasmon Resonance (SPR) on a Biacore T200 instrument using Protein-A coated S series chips. Antibodies were captured for 60 seconds at 5 µL/min to a level of 1000-2000RU onto the surface. The resurfaced RBDs were subsequently injected as analytes at five concentrations using the single cycle injection method, where no surface regeneration is performed between the analyte injections. The association phase was carried out for 180 seconds and the dissociation was done for 60 seconds with HBS-EP+ buffer flowing at 30 µL/min. Regeneration of the binding surface to capture a subsequent target antibody was done in 10 mM glycine-HC, pH 2 for 30 seconds at 30 µL/min with a 30 second baseline stabilization. A 1:1 Langmuir or Heterogenous ligand model was used for data fitting and analysis using the instrument software.

#### Antibody binding by ELISA

Enzyme-linked immunosorbent assays (ELISAs) were performed as described previously^12^. The following spike proteins were used, either produced as above or available commercially: WA-2 2P; SARS-CoV-1 2P; SHC014 2P; RaTG13 2P; and Gamma (Sino Biologicals, #40589-V08B10); Delta (Sino Biologicals, #40589-V08B16); BA.1.1 (Sino Biologicals, #40589-V08H29); BA.4/BA.5 (Sino Biologicals, #40589-V08H32); BQ.1.1 (Sino Biologicals, #40589-V08H41); XBB (Sino Biologicals, ref#40589-V08H40), Hexapro; XBB.1.5 (Sino Biologicals, #40589-V08H45); XBB.1.16 (Sino Biologicals, #40589-V08H49); EG.5.1 (Sino Biologicals, #40589-V08H55). Spikes and resurfaced RBDs were coated overnight on 384-well plates (Corning, 3700) at 4°C. Purified antibodies were used at 100 μg/mL starting concentrations and were then serially diluted 3-fold and incubated at room temperature for 1 hour. Goat anti-human IgG-HRP secondary antibody (Jackson ImmunoResearch Laboratories, 109-035-098) was diluted 1:15,000 and added to plates, incubated for one hour, and developed using TMB substrate (SureBlue Reserve, KPL, 5120-0083). Plates were read at 450 nm on a Cytation 1 plate reader (BioTek). Data was analyzed and plotted using GraphPad Prism version 10.2.0 for Windows (GraphPad Software, Boston, Massachusetts USA, www.graphpad.com). Statistics were performed by the Wilcoxon signed-rank test.

#### Sera binding by ELISA

ELISA binding assays with animal and human sera were performed as described previously^12^. Human sera was obtained from studies approved by the Duke University Health System Institutional Review Board (IRB) and conducted in agreement with the policies and protocols approved by the Duke IRB, consistent with the Declaration of Helsinki. Written informed consent was obtained from all research subjects or their legally authorized representatives. A full description of the cohort subjects and their SARS-CoV-2 exposure was provided previously^12^.

384-well plates were prepared in the same manner as in *Antibody binding by ELISA*. Human sera was diluted 1:30 and serially diluted 3-fold. Monomer mouse sera was diluted 1:100 while nanoparticle mouse sera was diluted 1:50 and serially diluted 5-fold. Samples were added and incubated at room temperature for 1.5 hours. The corresponding IgG-HRP secondary antibody was diluted and subsequently added (goat anti-human IgG-HRP, 1:15,000 dilution, Jackson ImmunoResearch Laboratories, 109-035-098; or goat anti-mouse IgG-HRP, 1:10,000 dilution, SouthernBiotech, 1030-05). After 1 hour incubation, plates were developed using TMB substrate (SureBlue Reserve, KPL, 5120-0083) and read at 450 nm on a SpectraMax Plus plate reader (Molecular Devices). Data was analyzed and plotted using GraphPad Prism version 10.2.0 for Windows (GraphPad Software, Boston, Massachusetts USA, www.graphpad.com). Statistics were performed by the Wilcoxon signed-rank test.

#### Sera blocking by ELISA

WA-2 spike was diluted to 2 μg/mL in 0.1 M Sodium Bicarbonate (Sigma-Aldrich, S6297) and coated overnight on 384-well plates at 4°C. Plates were washed with Superwash buffer (1X PBS supplemented with 0.1% Tween-20) and blocked for 1 hour at room temperature with Superblock buffer supplemented with azide (80 g Whey Protein, 300 mL Goat Serum, 20 mL aqueous 5% Sodium Azide, 10 mL Tween-20, 80 mL of 25X PBS, diluted to 2 L). Mouse sera was diluted 1:50 (in triplicate) in Superblock with azide. Samples were added and incubated at room temperature for 1.5 hours. In a separate dilution plate, biotinylated monoclonal antibodies were diluted at a final concentration equal to the half-maximal effective concentration at which they bound to spike. Plates were washed twice with Superwash, biotinylated mAbs were added, and allowed to incubate at room temperature for 1 hour. Plates were washed twice with Superwash and Strep-HRP secondary antibody (Streptavidin-HRP, 1:30,000 dilution, Thermo Scientific, 21130) in Superblock without azide was subsequently added. After incubating at room temperature for 1 hour, plates were washed four times in Superwash. Plates were developed with room temperature TMB substrate (SureBlue Reserve, KPL, 5120-0083) until the control wells reached an optical density (OD) of 1 and the reaction was stopped by 0. 33 M HCl solution. The OD at 450 nm was determined on a SpectraMax Plus plate reader (Molecular Devices). Data was analyzed and plotted using GraphPad Prism version 10.2.0 for Windows (GraphPad Software, Boston, Massachusetts USA, www.graphpad.com). Statistics were performed by the Wilcoxon signed-rank test.

### Mouse immunization studies

BALB/c female mice were purchased from Charles River. All mouse studies were performed under an approved Duke University IACUC protocol. All animal rooms were kept on a 12/12 light cycle unless otherwise requested. Heat and humidity were maintained within the parameters outlined in The Guide for the Care and Use of Laboratory Animals. Animals were fed a standard rodent diet. Mice were housed in individually ventilated micro-isolator caging on corn cob bedding. 3M052+Alumn in a 0.5+50 milligram mixture was used as the adjuvant for all the protein immunogens across the studies. Groups of mice *(n = 8)* were immunized intramuscularly with 2P stabilized D614 spike as mRNA (20 μg), monomeric WA-2 RBD (25 μg), monomeric resurfaced RBD immunogens (25 μg), or nanoparticles displaying wither the WA-2 RBD (10 μg) or the resurfaced RBDs (10 μg). Mice were immunized at week 0 (post-prime), week 4 (post-boost 1), and week 8 (post-boost 2). Blood samples were collected 7 days prior to immunization (pre-bleed) and 7 days after each immunization. Mice were anesthetized in an induction box using isoflurane (5%) and were unconscious during each injection and blood collection. Eye drops were applied to each mouse to prevent the eyes from drying out during anesthesia. All mice received a total immunization volume of 50 μL via two intramuscular 25 μL injections into each bicep femoris using a 28-gauge sheathed needle. Seven days after the final dose animals were sacrificed for terminal analysis. For euthanasia, mice were first anesthetized in an induction box (5% isoflurane) for terminal blood collection. The mice were then transferred into a CO_2_ induction box in accordance with IACUC’s euthanasia protocol (4 L/min flow rate) until cessation of breathing. A secondary method of death was confirmed by bilateral thoracotomy.

### Sera neutralization

#### Live virus neutralization

To determine neutralization potential of antibodies in serum, an authentic SARS-CoV-2 microneutralization assay was used. Serum samples were heat inactivated at 56°C for one hour and then diluted two-fold in virus diluent (MEM + Earl’s Salts + L-glutaime (Gibco 11095), with 1X Penicillin/Streptomycin (Gibco 15140), 2% heat inactivated fetal bovine serum (Gemini 100-106), 1 mM sodium pyruvate (Gibco 11360), 1X MEM nonessential amino acids (Gibco 11140) and 10 μg/mL Puromycin (ThermoFiser A1113803). 100 TCID_50_ of the indicated virus strains was added to each well and incubated for 1 hour at 37°C. The serum-virus mixture was then transferred to wells containing 25,000 Vero E6 cells expressing TMPRSS2 and ACE2 (BEI resources NR-54970) that were prepared the day before infection. The cells were incubated for 19 hours at 37°C, fixed with 10% neutral buffered formalin for at least 30 minutes. Cells were made permeable with a Tween-20 buffer (ThermoFisher J62844.K3) supplemented with 0.01 M Glycine (Sigma G2879). Primary antibody (ThermoFisher MA5-29981) was used to detect SARS-CoV-2 nucleocapsid inside the cells, and a secondary antibody (Abcam ab150113) was used to generate fluorescent signal. Signal was captured by an automated plate reader, and data were expressed as the Mean Effective Concentration which is the reciprocal of the last dilution of serum to neutralize 50% of the virus specific signal. All samples were run in duplicate. All groups *n = 8* represent independent mice, except for mRNA + Resurf6 mi03-NP + Resurf61 mi03-NP *(n = 7)* due to a mouse death prior to terminal bleeds. Significance tested using the Wilcoxon signed-rank test.

#### Pseudovirus neutralization

Serum neutralization of pseudotyped coronavirus infection of ACE2-expressing 293T cells was evaluated. Serum aliquots were inactivated by heating at 56°C for 15 minutes in a water bath. Mouse serum was diluted in complete growth media (DMEM +10% FBS +1% Penicillin/Streptomycin) at a starting dilution of 1:80 (SARS-CoV-2 XBB.1 and SARS-CoV-JN.1) or 1:100 (SARS-CoV-2 D614G, SARS-CoV-1-Urbani and RsSHC014) in a total volume of 125 μL in a 96-well cell culture plate. Five-fold serial dilutions (25 μL) were performed 7 times in the same plate, and the final 25 μL was discarded. Fifty microliters of the tested pseudoviruses were added to each well at a predetermined dilution based on pseudovirus titration assays. Wells with complete growth media instead of serum were included for comparison. Plates were then incubated at 37°C with 5% CO_2_ for 1 hour. A suspension of 293T/ACE2 cells (kindly provided by Mike Farzan at The Scripps Institute) were prepared at 100,000 cells per milliliter per standard cell preparation techniques. After the 1-hour incubation between pseudotyped virus and serum, 100 μL of cell suspension (10,000 cells total) were added to each well. Plates were rested at room temperature for 30 minutes and then incubated at 37°C with 5% CO_2_ for 72 hours. After incubation, the entire media content of each well was aspirated and discarded, and 115 μL Promega BrightGlo Reagent was added and incubated in the plate in the dark for 2 minutes 30 seconds. The contents of each well were mixed with a multichannel pipette and 100 μL of the BrightGlo reagent was moved to the corresponding well of a new PerkinElmer Black/White microplate and imaged on a VictorNivo Luminometer. Data were analyzed via the Nab tool in the CHAVI Specman application and figures were created in GraphPad PRISM 10. All groups *n = 8* represent independent mice, except for mRNA + WA-1 RBD x2 *(n = 5*)* and mRNA + Resurf6 mi03-NP + Resurf61 mi03-NP *(n = 6**)*. *Sera from three mice were not available to be tested. **Sera from one mouse was not available to be tested and a mouse death occurred prior to terminal bleeds. Statistics were performed by the Mixed-effects model.

### Conservation Analysis

To map conservation of sarbecovirus sequences on to the RBD structure, sarbecovirus protein sequences were obtained from two distinct sources: NCBI BLAST^67^ and GISAID^68^. For sequences retrieved via BLAST, we utilized the BLAST command line application (version 2.12.0) and the non-redundant (nr) database (downloaded on January 17, 2024). The blastp (protein-protein) command was used to acquire accession numbers, utilizing a query sequence and a taxonomy ID list as input, with the e-value option set to 1 and max_target_seqs set to 10,000,000 to ensure all possible hits were returned. Two separate runs were conducted using different query sequences and taxonomy ID lists. The first run used the SARS-CoV-2 Omicron JN.1 spike protein from GISAID (EPI_ISL_18300149) as the query sequence and the taxonomy ID for SARS-CoV-2 (NCBI:txid2697049). The second run used the Bat SARS-like coronavirus RsSHC014 (KC881005.1:21492-25262) spike protein as the query sequence and all sarbecovirus taxonomy IDs with the SARS-CoV-2 taxonomy ID excluded. The taxonomy ID list of all sarbecoviruses was obtained using the get_species_taxids.sh script from the BLAST command line application with the sarbecovirus taxonomy ID as input (NCBI:txid2509511) and manually removing the SARS-CoV-2 taxonomy ID.

Following blastp, the dbcmd tool from BLAST was employed to extract the full-length sequences using the accession numbers from the blastp output for both runs. Subsequently, a custom semi-global alignment script was utilized to extract the spike regions from both sets of sequences, using the previously described query sequences as references. Finally, the RBD regions were extracted from the spikes using the same custom semi-global alignment script with the respective RBD region from each query sequence as reference.

For sequences retrieved from GISAID, SARS-CoV-2 spikes available on March 17, 2024, were downloaded. Sequences from non-human hosts were then excluded. The remaining sequences were deduplicated based on sequence i.e. if two or more sequences where identical, one was kept and the rest removed to retain only unique sequences. The same semi-global alignment script was then utilized to extract the RBD region using the Wuhan WIV04 RBD sequence (EPI_ISL_402124) as a reference. Finally, all SARS-CoV-2 sequences were combined.

To include only high-quality samples in this analysis, sequences were filtered to retain those with lengths between 220 and 226 amino acids and containing fewer than two gap (’X’) characters. The length criteria was determined according to the typical length of SARS-CoV-2 RBDs, which are generally longer than those of other sarbecoviruses^69^. This criteria was crucial to eliminate low-quality sequences often characterized by large gaps, as a result from sampling late in infection, which yields low viral loads. Such sequences could also arise from technical challenges during sample preparation, sequencing, and bioinformatics processing^70^. A total of 2,091,113 unique SARS-CoV-2 sequences meeting these criteria were obtained.

Following filtering, a hierarchical clustering approach was employed to obtain a representative set of sequences covering the diversity of SARS-CoV-2. First, sequences were clustered at 92% identity using the cluster_fast command in USEARCHs (v10.0.240)^71^, resulting in 44 clusters, with sizes ranging from 1 to 1,566,099. Except for clusters containing a single sequence, each cluster was further clustered at 96% identity, with the sort option set to size to improve clustering efficiency. The largest number of subclusters produced was 131. To ensure a balanced representation, the number of sequences from each cluster was standardized to the maximum number of subclusters obtained, 131. For smaller clusters, centroids were sampled with replacement to match this number, preventing oversampling and overrepresentation in the final sequence set. This process resulted in a total of 5,764 sequences.

A similar approach was applied to non-SARS-CoV-2 sarbecovirus sequences (n = 2,088,366), with a relaxed length threshold of ≥ 200 residues to include clade 2 sarbecoviruses, which have shorter RBDs (204-205 residues)^72^. Clustering at 92% identity produced 66 clusters, with the largest subcluster containing 1,564,379 sequences. The final sequence sets included 8,052 non-SARs-CoV-2 sarbecovirus sequences and 5,453 SARS-CoV-2 sequences.

The sequences were then aligned using MAFFT^73^. Conservation scores for each position in the multiple sequence alignment were calculated using the trident scoring method^74^ implemented in the MstatX program (https://github.com/gcollet/MstatX). The conservation scores were then mapped to the domain coordinates of the SARS-CoV-2 RBD (PDB: 6LZG) B-factor column^75^. A custom Perl script was used to obtain the binding footprint of the antibody in complex with the RBD and images rendered with PyMol version 3.0.0^76–78^.

Data analysis and visualization was performed in the R-programming language. Group medians were compared using exact Wilcoxon test using the exactRankTests R package. P values of ≤ 0.05, ≤ 0.01, and ≤ 0.001 determined increased significance levels.

## Data availability

Any materials and data described in this study will be made available in a timely fashion to members of the scientific community for noncommercial purposes.

## Acknowledgments

We thank Whitney Edwards Beck for program management help and for assistance with acquiring and managing the human samples. We thank Brian Watts, Ken Cronin, and Alam Munir for their assistance at the DHVI Biomolecular Interaction Analysis (BIA) Core Facility. We thank Wes Rountree and Yunfei Wang for the statistical analysis. We thank Jacob Gater and Pahvie Chhan from the Duke Human Vaccine Institute Small Animal Team for their help with the immunization studies. This work was supported by grants P01AI158571 (B.F.H) and R01AI155804 (M.L.A) from the National Institute of Allergy and Infectious Diseases of the National Institute of Health and by a Duke School of Medicine COVID Award (M.L.A).

## Author contribution

ABK and MLA designed the study and the resurfaced RBDs; ABK and COD produced the engineered resurfaced RBDs; CH performed all the sera and antibody binding assays; CH, RP and MB performed sera blocking assays; JC and JL performed melting experiments of the resurfaced RBDs; JES, MB and KW performed the spike sequence conservation analysis; KJ, SS, XH, JL and PA provided spike and RBD proteins for sera and antibody analysis; TO performed live virus neutralization assays; TG, MO, and KOS performed pseudovirus neutralization; KOS provided the ferritin-WT RBD nanoparticle; AN performed the mouse immunizations; PA, BFH, KW, KOS and MLA provided project supervision; BFH and MLA secured the funding; CH and JES generated the figures with input from KW and MLA; MLA ABK, CH, JES and MLA wrote the first draft on which all authors commented.

