## Supplementary Figures for "Design of SARS-CoV-2 RBD Immunogens to Focus Immune Responses Towards Conserved Coronavirus Epitopes"

**Binding Site Conservation Scores**

| Position | S309 |
|---|---|
| 333 | 0.968 |
| 334 | 0.968 |
| 335 | 0.875 |
| 336 | 0.981 |
| 337 | 0.963 |
| 339 | 0.662 |
| 340 | 0.822 |
| 341 | 0.974 |
| 356 | 0.804 |
| 358 | 0.958 |
| 359 | 0.934 |
| 360 | 0.837 |
| 361 | 0.909 |
| 441 | 0.711 |
| 505 | 0.806 |
| 509 | 0.977 |

| Position | S2X259 |
|---|---|
| 370 | 0.979 |
| 371 | 0.706 |
| 372 | 0.721 |
| 373 | 0.722 |
| 374 | 0.943 |
| 375 | 0.757 |
| 376 | 0.807 |
| 377 | 0.953 |
| 378 | 0.883 |
| 379 | 0.945 |
| 380 | 0.934 |
| 381 | 0.951 |
| 383 | 0.928 |
| 384 | 0.821 |
| 385 | 0.804 |
| 405 | 0.719 |
| 408 | 0.789 |
| 502 | 0.826 |
| 503 | 0.704 |
| 504 | 0.794 |
| 506 | 1.000 |
| 507 | 0.892 |

| Position | ACE2 |
|---|---|
| 403 | 0.817 |
| 445 | 0.700 |
| 446 | 0.682 |
| 449 | 0.755 |
| 453 | 0.965 |
| 455 | 0.746 |
| 456 | 0.758 |
| 473 | 0.735 |
| 475 | 0.714 |
| 476 | 0.676 |
| 478 | 0.615 |
| 484 | 0.462 |
| 485 | 0.712 |
| 486 | 0.647 |
| 487 | 0.886 |
| 489 | 0.875 |
| 490 | 0.638 |
| 493 | 0.659 |
| 494 | 0.727 |
| 495 | 0.947 |
| 496 | 0.780 |
| 497 | 0.904 |
| 498 | 0.525 |
| 501 | 0.535 |

**Figure S1**

**Binding site conservation scores across the individual RBD amino acids involved in interactions with S309 mAb (*orange*), S2X259 mAb (*pink*), and ACE2 (*grey*), respectively.**

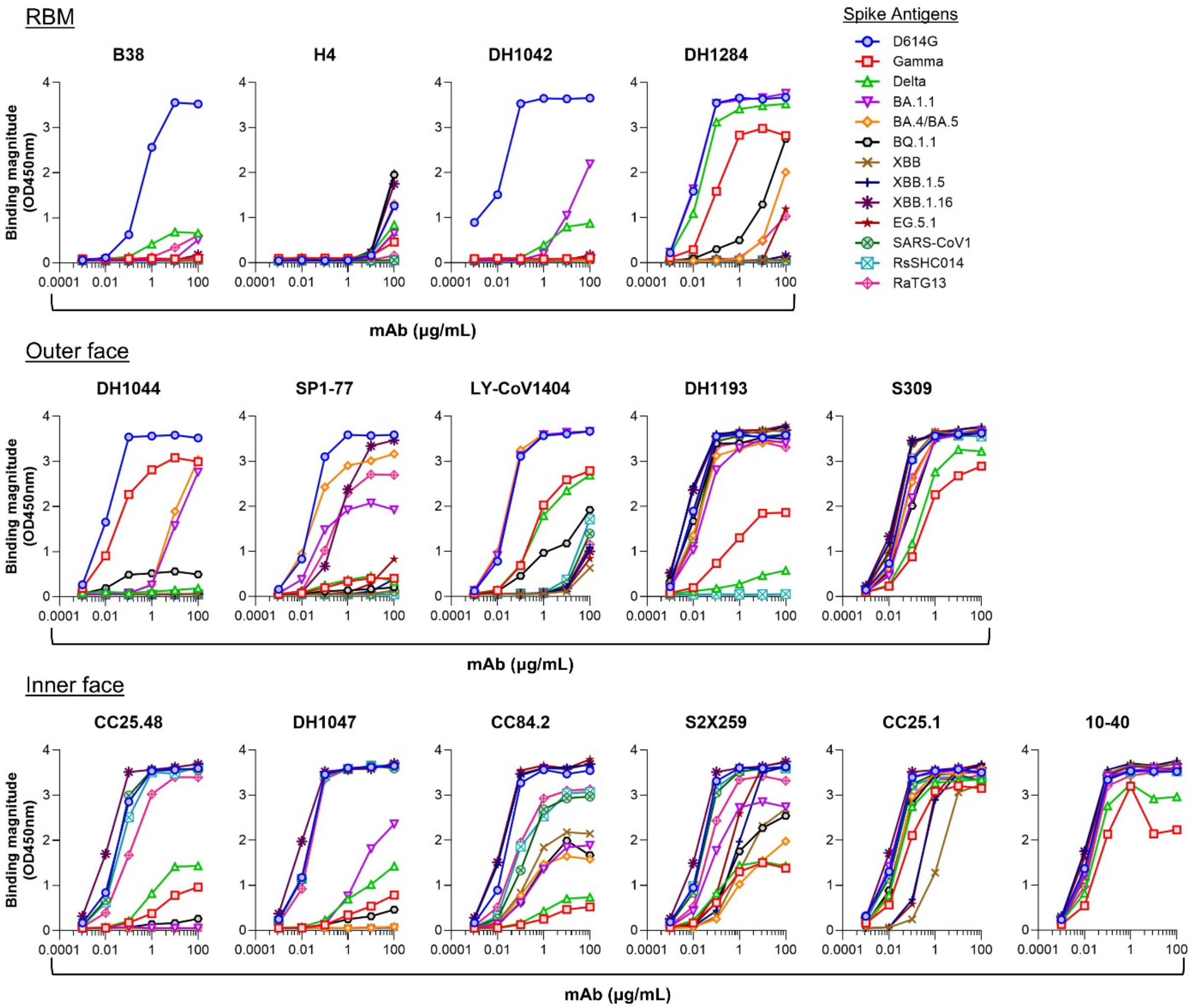


**Figure S2**

**ELISA binding titration curves of RBM, outer and inner face antibodies to SARS-CoV-2 VoC spikes.**

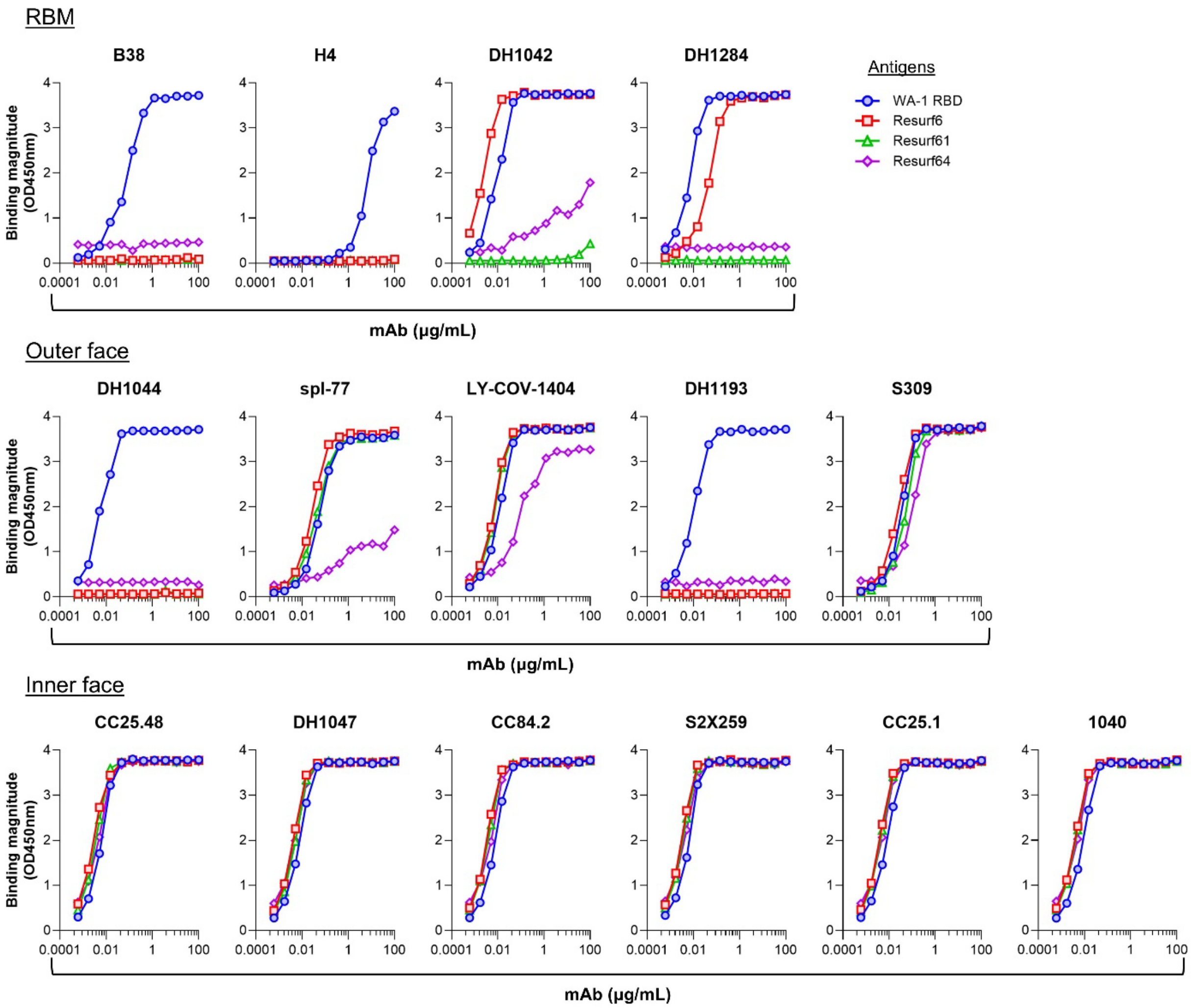


**Figure S3**

**ELISA binding titration curves of RBM, outer and inner face antibodies to WA-1 RBD and resurfaced RBD immunogens.**

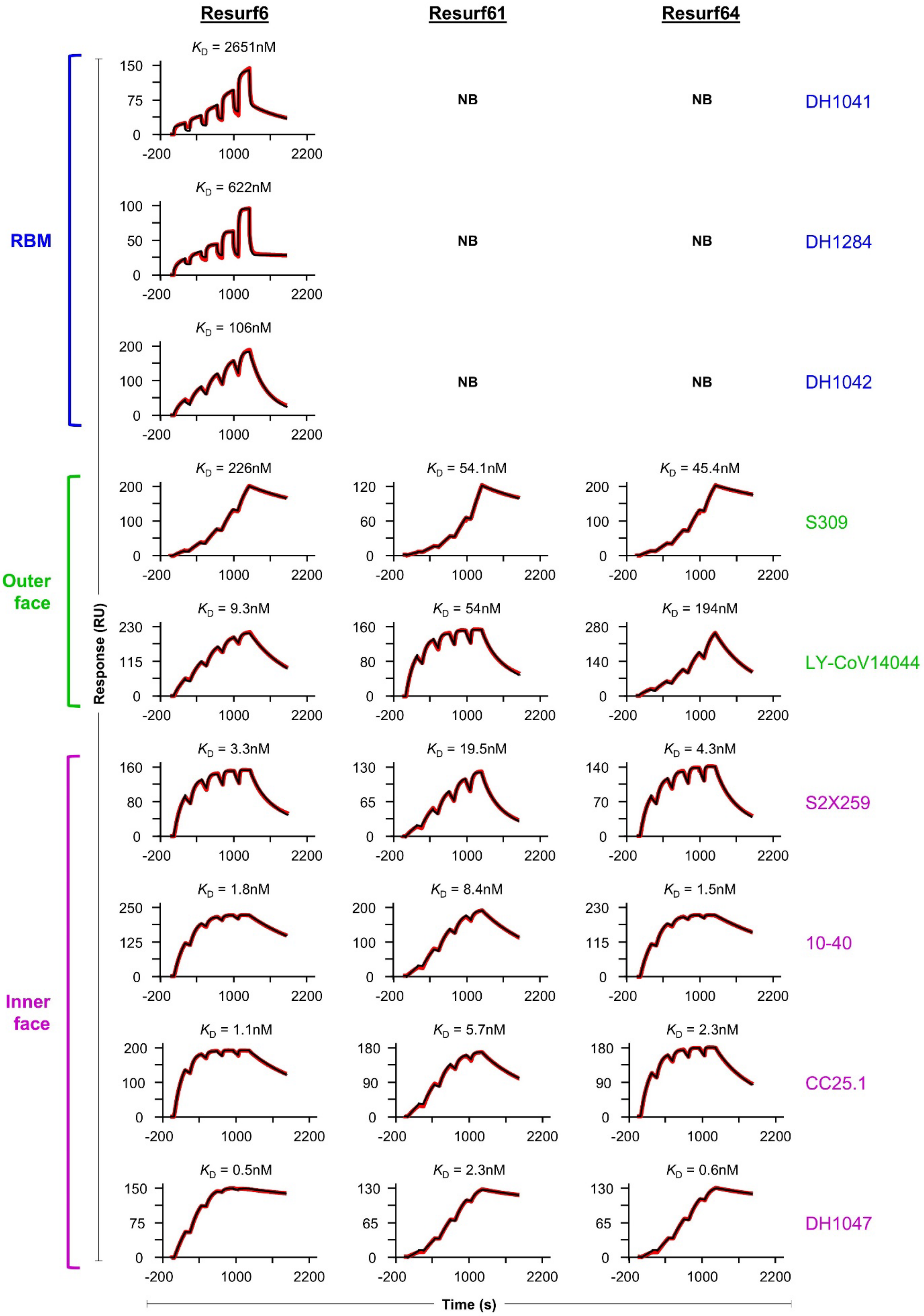

Resurf6
Resurf61
Resurf64
RBM
Outer face
Inner face
Response (RU)
Time (s)
DH1041
DH1284
DH1042
S309
LY-CoV14044
S2X259
10-40
CC25.1
DH1047
$K_D$ = 2651nM
$K_D$ = 622nM
$K_D$ = 106nM
NB
$K_D$ = 226nM
$K_D$ = 54.1nM
$K_D$ = 45.4nM
$K_D$ = 9.3nM
$K_D$ = 54nM
$K_D$ = 194nM
$K_D$ = 3.3nM
$K_D$ = 19.5nM
$K_D$ = 4.3nM
$K_D$ = 1.8nM
$K_D$ = 8.4nM
$K_D$ = 1.5nM
$K_D$ = 1.1nM
$K_D$ = 5.7nM
$K_D$ = 2.3nM
$K_D$ = 0.5nM
$K_D$ = 2.3nM
$K_D$ = 0.6nM

**Figure S4**

**Binding affinities of resurfaced RBD immunogens to RBM, outer and inner antibodies.** Single cycle Surface Plasmon Resonance (SPR) kinetic measurements of Resurf6, Resurf61, and Resurf64 binding to DH1041, DH1284, DH1042, S309, LY-CoV1404, S2X259, 10-40, CC25.1, and DH1047 mAbs.

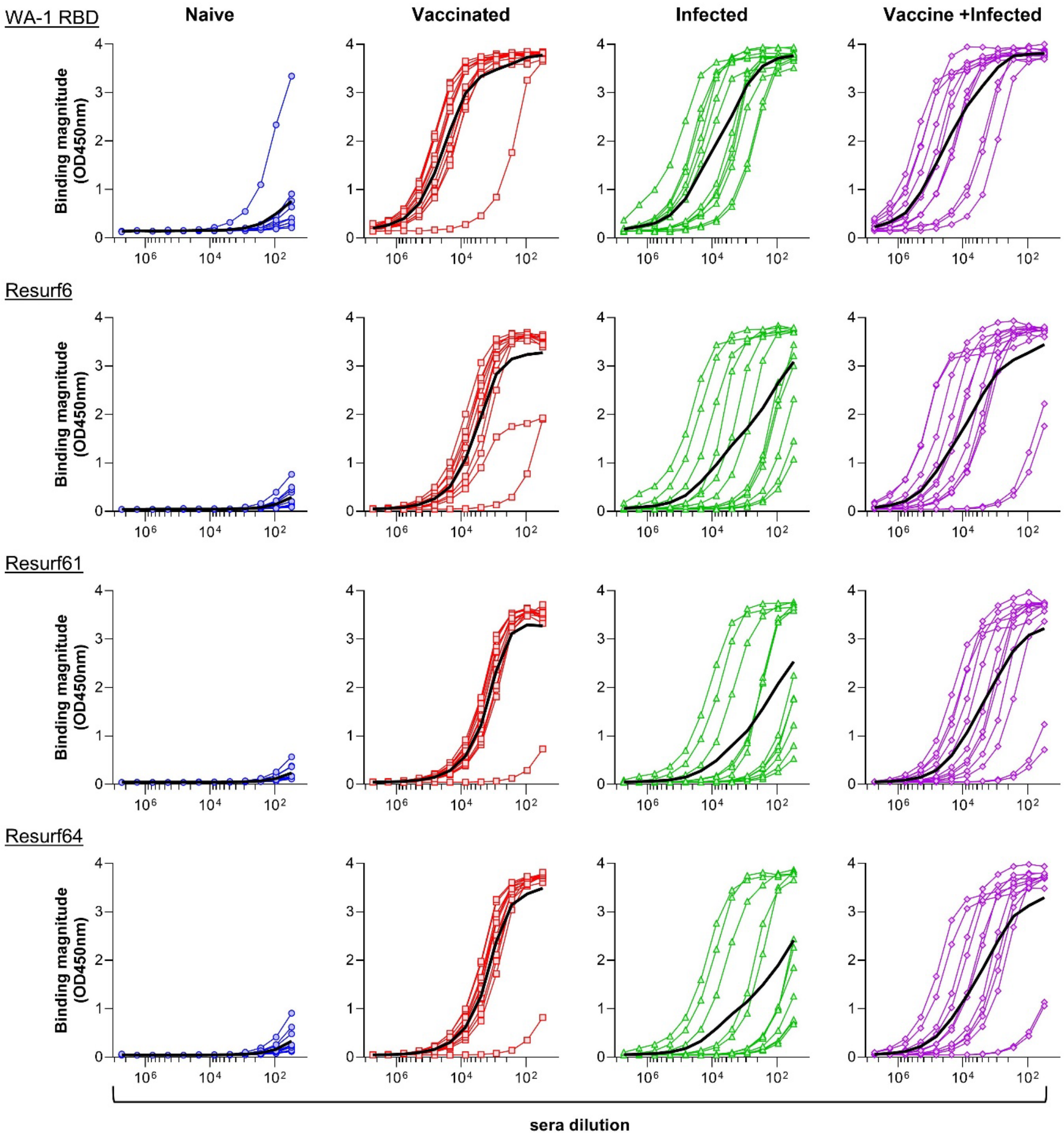


**Figure S5**

**Binding of human sera from subjects naïve (*blue*) or with existing SARS-CoV-2 immunity acquired by vaccination (*red*), infection (*green*), or vaccination followed by infection (*purple*) to WA-1 RBD and engineered RBDs.** Lines represent individual measurements for each subject in the group *(n = 12)*. Group averages in *black* lines.

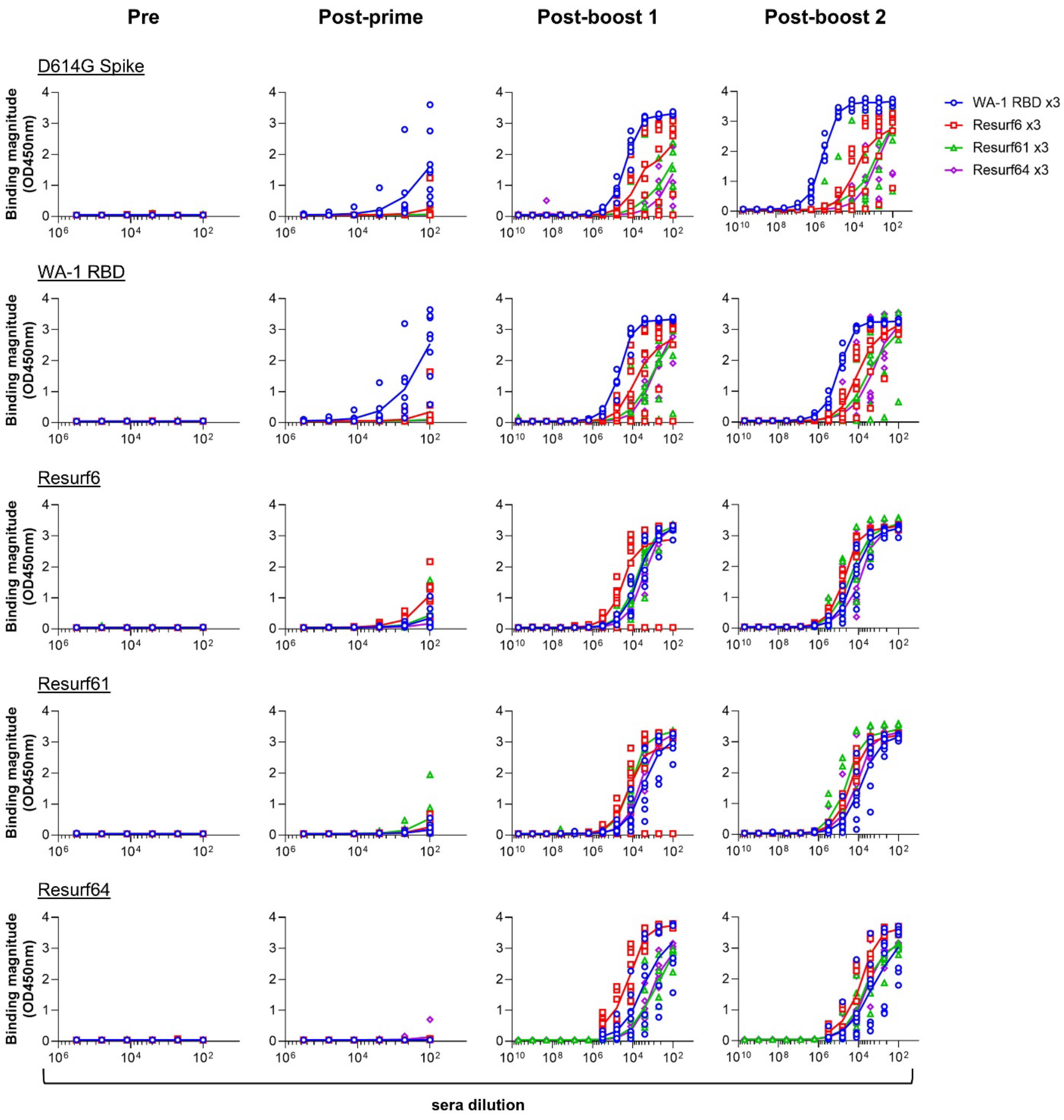


**Figure S6**

**ELISA binding titration curves of sera from BALB/c mice immunized with WA-1 RBD or resurfaced immunogens to D614G spike, WA-1 RBD, and resurfaced RBD immunogens.** D614G spike = SARS-CoV-2 variant. Sera binding at pre-prime, post-prime, post-boost 1, and post-boost 2 time points *(n = 8)*.

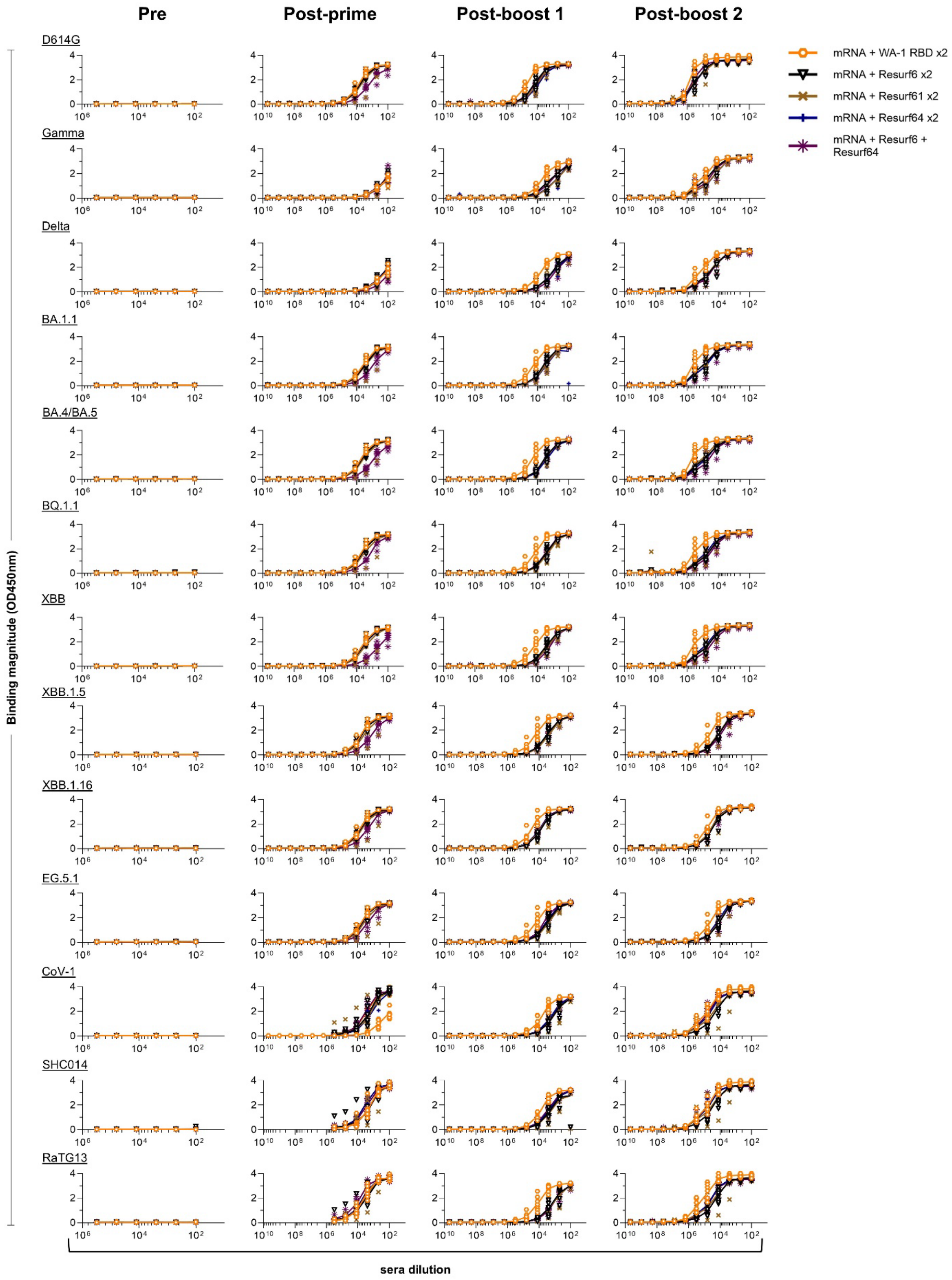
Pre
Post-prime
Post-boost 1
Post-boost 2
D614G
Gamma
Delta
BA.1.1
BA.4/BA.5
BQ.1.1
XBB
XBB.1.5
XBB.1.16
EG.5.1
CoV-1
SHC014
RaTG13
Binding magnitude (OD450nm)
sera dilution
mRNA + WA-1 RBD x2
mRNA + Resurf6 x2
mRNA + Resurf61 x2
mRNA + Resurf64 x2
mRNA + Resurf6 + Resurf64

**Figure S7**

**ELISA binding titration curves of sera from mice primed with mRNA and boosted twice with different RBD immunogen to diverse CoV spikes.** D614G, Gamma, Delta, BA.1.1, BA.4/BA.5, BQ.1.1, XBB, XBB.1.5, XBB.1.16 and EG.5.1 refer to the SARS-CoV-2 variants. CoV-1 = SARS-CoV-1; SHC014 = BatCoV RsSHC014; RaTG13 = BatCoV RaTG13. Sera binding at pre-prime, post-prime, post-boost 1, and post-boost 2 time points *(n = 8)*.

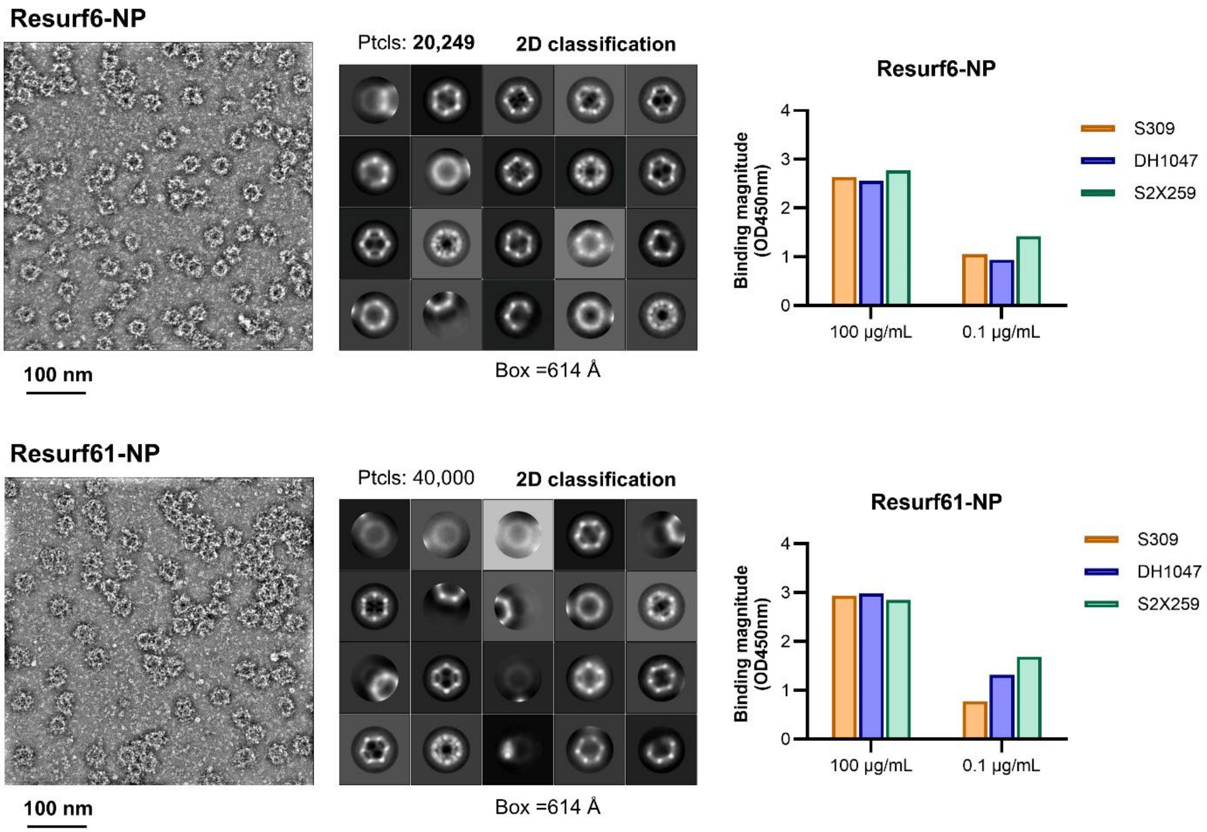


**Figure S8**

***In vitro* characterization of resurfaced RBD nanoparticles.** *Left*: NSEM analysis of resurfaced RBDs multimerized on mi03 nanoparticles. *Right*: Antigenicity of resurfaced RBD nanoparticles.

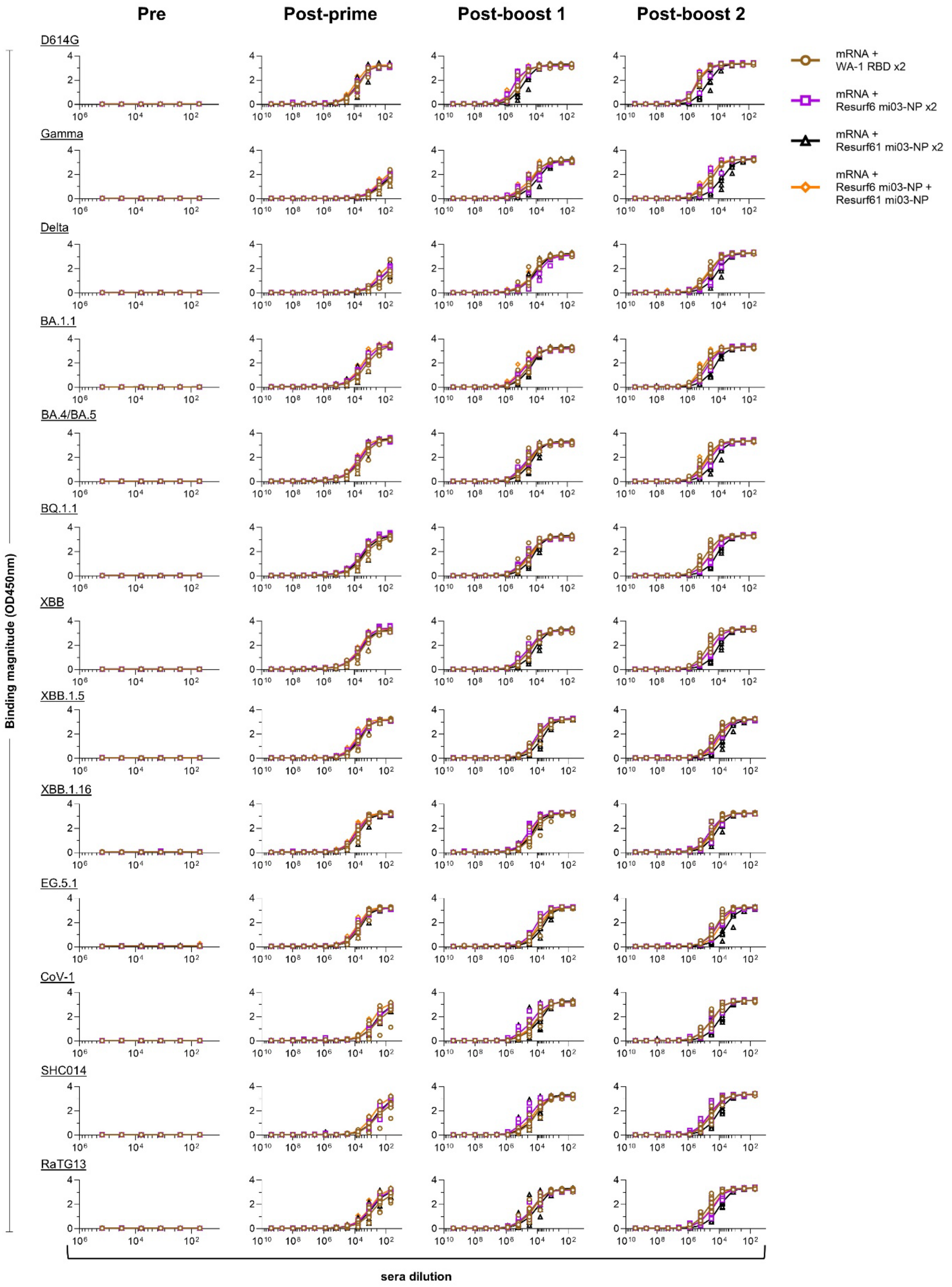

Pre
Post-prime
Post-boost 1
Post-boost 2
D614G
Gamma
Delta
BA.1.1
BA.4/BA.5
BQ.1.1
XBB
XBB.1.5
XBB.1.16
EG.5.1
CoV-1
SHC014
RaTG13
Binding magnitude (OD450nm)
sera dilution
mRNA + WA-1 RBD x2
mRNA + Resurf6 mi03-NP x2
mRNA + Resurf61 mi03-NP x2
mRNA + Resurf6 mi03-NP + Resurf61 mi03-NP


**Figure S9**

**ELISA binding titration curves of sera from mice primed with mRNA and boosted twice with different RBD nanoparticle (NP) immunogen to diverse CoV spikes.** D614G, Gamma, Delta, BA.1.1, BA.4/BA.5, BQ.1.1, XBB, XBB.1.5, XBB.1.16 and EG.5.1 refer to the SARS-CoV-2 variants. CoV-1 = SARS-CoV-1; SHC014 = BatCoV RsSHC014; RaTG13 = BatCoV RaTG13. Sera binding at pre-prime, post-prime, post-boost 1, and post-boost 2 time points *(n = 8)*.

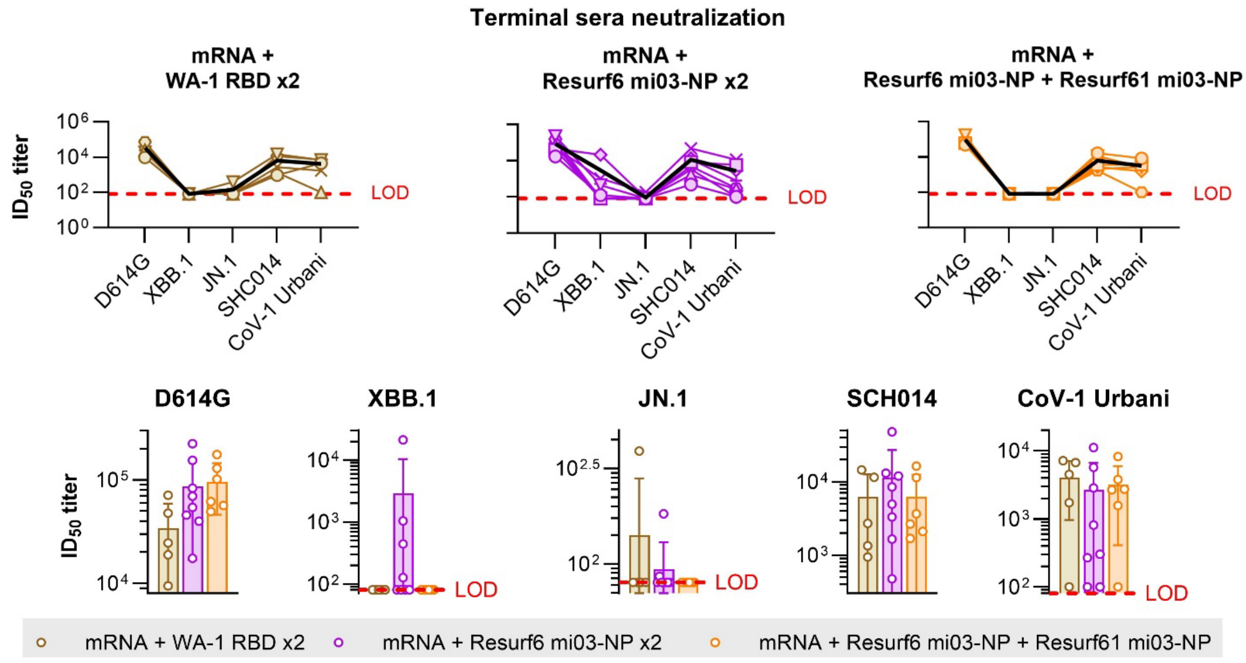


### Figure S10

**Pseudovirus neutralization of terminal sera from animals primed with mRNA spike and boosted with RBD immunogen nanoparticles.** *Top*: data plotted for individual animals in each group against the pseudovirus panel. Group averages in *black* lines. Level of detection (LOD) in *red*. *Bottom:* average group neutralization titers against each pseudovirus. All groups *n = 8* represent independent mice, except for mRNA + WA-1 RBD x2 *(n = 5*)* and mRNA + Resurf6 mi03-NP + Resurf61 mi03-NP *(n = 6**)*. *Sera from three mice were not available to be tested. **Sera from one mouse was not available to be tested and a mouse death occurred prior to terminal bleeds. Error bars show +/- standard deviation. Statistics were performed by the Mixed-effects model.
